# Dissecting signalling hierarchies in the patterning of the mouse primitive streak using micro-patterned EpiLC colonies

**DOI:** 10.1101/2020.11.30.404418

**Authors:** Jean-Louis Plouhinec, Mathieu Vieira, Gaël Simon, Jérôme Collignon, Benoit Sorre

**Affiliations:** Université de Paris, CNRS, Laboratoire Matière et système Complexes, Paris France; Université de Paris, CNRS, Institut Jacques Monod, Paris, France

## Abstract

Molecular embryology studies have established that the patterning of the gastrula-stage mouse embryo is dependent on a regulatory network where the WNT, BMP and NODAL signalling pathways cooperate. Still, important aspects of their respective contributions to this process remain unclear. Here, studying their impact on the spatial organization and the developmental trajectories of micro-patterned Epiblast Like Cells (EpiLC) colonies, we show that when BMP is present, it dominates NODAL and WNT and imposes a posterior character to the colonies differentiation. However, the use of two *Nodal* mutant cell lines allowed us to show that prior to BMP action, NODAL is required to establish the mesendodermal lineage. The fact that mutant phenotypes were more severe *in vitro* than *in vivo* suggests that embryonic phenotypes are partially rescued by ligands of extra-embryonic or maternal origin. Our work demonstrates the complementarity of micro-patterned EpiLC colonies to embryological approaches.

## Introduction

The epiblast is the pluripotent embryonic tissue that gives rise to the three germ layers - the ectoderm, the mesoderm and the endoderm - which then go on to form all foetal tissues. Theoretical work suggested that such tissue patterning must involve signalling molecules, providing naïve undifferentiated cells with the means to assess either their position in the tissue or the status of neighbouring cells, and to determine their fate accordingly (Meinhardt, Hans, 1982; Wolpert, 1969). Mouse genetics has indeed confirmed or revealed the implication of a number of ligands and signal transducers, notably for the WNT/β-CATENIN, BMP, and ACTIVIN/NODAL signalling pathways, in the patterning of the epiblast (Arnold and Robertson, 2009; Tam and Loebel, 2007). This process culminates in the formation of the primitive streak (PS), the structure through which posterior epiblast cells ingress as they undergo the epithelial to mesenchymal transition (EMT) associated with the adoption of mesendodermal cell identities. The PS itself is patterned; cells emerging at different times and at different levels of the PS will contribute to distinct embryonic regions and will adopt fates appropriate for their positions. Cells emerging early on from the proximal/posterior region of the PS will thus form extra-embryonic mesoderm and embryonic mesoderm while cells emerging from its distal/anterior region a few hours later will form axial mesoderm and definitive endoderm (Kinder et al., 1999).

Three ligands, WNT3, BMP4 and NODAL, were found to be critical for the formation and the patterning of the PS. Loss-of-function mutations in either of the corresponding genes appeared to result in the absence of a PS (Conlon et al., 1991, 1994; Iannaccone et al., 1992; Liu et al., 1999; Winnier et al., 1995; Zhou et al., 1993). However, anterior mesoderm cells were found in some *Bmp4^-/-^* embryos and posterior mesoderm markers were found in some *Nodal^-/-^* embryos (Ben-Haim et al., 2006; Conlon et al., 1994), suggesting distinct roles in PS patterning. The study of an hypomorphic allele of *Nodal* and of mutant alleles of its transducers *Smad2* and *Smad3* implied that cell fate decisions in the streak are dependent on the strength of this signal, its gradual reduction resulting first in the disappearance of the most anterior mesendodermal fates, including the definitive endoderm, and then in the additional disappearance of more posterior mesodermal fates (Vincent et al., 2003). These results were consistent with others obtained in *Xenopus*, which showed that BMP4 promotes the differentiation of posterior mesoderm and counteracts the opposing effect of dorsalizing factors such as ACTIVINs and NODAL, which act in concert with the WNT/β-CATENIN pathway (Harland, 1994; Zorn et al., 1999). They are at the basis of current models, where opposing gradients of NODAL and BMP4 signalling activities govern the A/P patterning of PS derivatives in the mouse (Arnold and Robertson, 2009; Tam and Loebel, 2007).

While developmental genetics is apparently consistent with cell fate allocation in the mouse PS being dependent on the level of NODAL signalling, and thus reflecting its gradation along the A/P axis, there is actually no clear evidence of such gradation. Detection of *Nodal* transcripts via *in situ* hybridization and a reporter transgene for NODAL signalling both revealed broadly homogeneous expression patterns along the A/P axis at relevant stages (Norris et al., 2002; Robertson et al., 2003), suggesting further complexity. One possibility is that cell fate allocation reflects how long cells have been exposed to NODAL during their transit through the PS. Testing such hypotheses in the developing mouse embryo is at best challenging, sometimes impossible.

Indeed, one difficulty when studying how the epiblast and the PS are actually patterned stems from the demonstrated interdependency of *Wnt3, Bmp4* and *Nodal. Nodal* expression in the epiblast promotes that of *Bmp4* in the adjacent extra-embryonic ectoderm, which promotes the expression of *Wnt3* in the posterior epiblast, which in turn amplifies *Nodal* expression there via an enhancer dependent on canonical WNT signalling (Ben-Haim et al., 2006; Granier et al., 2011). This interdependency and the fact that these genes also have earlier roles in embryo patterning make it particularly challenging to design experiments to distinguish, for example, the effect of the lack of NODAL from the effect of a lack of BMP4 and WNT3 in the patterning defects of *Nodal^-/-^* embryos.

The recent development of methods that use pluripotent cells to recapitulate key aspects of gastrulation *in vitro*, thus allowing greater control over the conditions under which patterning takes place, now provides possible alternatives for such investigations. In particular, human embryonic stem cells (hESCs) cultured as a monolayer on adhesive circular micropatterns, similar in size to that of an embryo, were found to self-organize when exposed to BMP4, and to give rise to the three embryonic germ layers arranged in concentric rings in an ordered and reproducible sequence (Warmflash et al., 2014). This 2D *in vitro* model of patterning, or 2D gastruloid, has many advantages. It is simple, multiple colonies can be generated at once and it is amenable to imaging and quantification. However, there is evidence of notable differences between human and murine development (Blakeley et al., 2015; Etoc et al., 2016), and to allow the comparison of the events taking place *in vitro* with those actually taking place in the embryo we needed to adapt this method to the use of mouse pluripotent stem cells.

Reporting on this undertaking, we confirm that EpiLCs, pluripotent cells derived from mouse ESCs (mESCs), can generate a reproducible differentiation pattern when induced by BMP4 (Morgani et al., 2018). We used RT-PCR and principal component analysis (PCA) to efficiently characterize the developmental trajectories of differentiating colonies and to compare the inducing capabilities of BMP4, ACTIVIN+FGF2 (a proxy for NODAL) and WNT3a (a proxy for WNT3). In line with the results of embryological studies, we found that BMP4 promotes the formation of proximal/posterior cell identities while WNT3a promotes the formation of distal/anterior ones. Furthermore, we found that sustained exposure to BMP4 prevents the emergence of anterior fates, regardless of the addition of other signalling molecules. However, a partial and a complete depletion of NODAL, obtained via the use of two distinct mutant *Nodal* alleles, had more severe effects on colony patterning than what previous studies of mutant embryonic phenotypes led us to expect. The defects were consistent with *Nodal* being required first to endow epiblast cells with the capacity to form a PS, and then to promote the formation of anterior PS derivatives. The severity of the *in vitro* phenotypes hints at the possibility that the embryonic phenotypes are partially rescued by ligands of extra-embryonic or maternal origin. Our study thus delineates an efficient approach for the study of embryo patterning *in vitro* and demonstrates its complementarity to embryological approaches.

## Results

### 1. 2D-patterning of mouse pluripotent stem cell colonies

Different mouse pluripotent stem cell populations were considered to model in culture the maturation and differentiation of the epiblast taking place in the embryo in response to inductive cues. ESCs are derived from the preimplantation epiblast and are in a ground state of pluripotency. They express a combination of pluripotency factors that is distinctive of their identity and they are not readily responsive to inductive signals in conditions allowing them to self-renew. EpiSCs (Epiblast stem cells) are derived from post-implantation epiblast and are in a state of primed pluripotency (Brons et al., 2007; Tesar et al., 2007). They do not express some of the pluripotency factors seen in ESCs but they express lineage-specific markers and they show heterogeneity in their differentiation capabilities (Sugimoto et al., 2015). EpiLCs (EpiSC-like cells) are obtained when ESCs are cultured for 2-3 days in the presence of FGF and ACTIVIN (Hayashi et al., 2011). They are transcriptionally distinct from both ESCs and EpiSCs, and closer to early post-implantation epiblast (Hayashi et al., 2011; Kalkan et al., 2017). They have lost pluripotency factors that are distinctive of ESCs but do not express regionalized lineage markers. They are homogeneous and have acquired the capacity to respond to inductive signals and are thus said to be in a state of formative pluripotency (Smith, 2017). They were the natural choice when attempting to recapitulate *in vitro* the patterning events associated with gastrulation.

To initiate their conversion into EpiLCs, mESCs were seeded on Fibronectin-coated petri dishes in N2B27+ACTIVIN+FGF medium (t=-48h, Fig. 1A). After 24 hours, cells were trypsinized, seeded on adhesive micro-patterned substrates (700μm diameter) obtained by micro-contact printing of Fibronectin on PDMS-coated glass slides (see online methods for a detailed protocol), and allowed to spread on these substrates for another 24h. This 2-step protocol ensures the homogeneous seeding of adhesive micro-patterns, which is necessary to obtain reproducible patterning of the colonies. 48h after the start of the culture (t=0, Fig. 1A), cells in the colonies had an expression profile consistent with the acquisition of an EpiLC identity - spatially homogenous expression of OCT4 and E-CAD (Fig. S1A), downregulation of *Sox2* and *Nanog* expression, gain of *Otx2* and *Fgf5* expression (Fig. S1B) - indicating that they were ready to be stimulated. BMP4 (50ng/ml) was thus added to the EpiLC differentiation medium, and the spatial organization of cell identities in the colonies was characterized by immunofluorescence (IF) after 24, 48 and 72h of culture.

**Figure 1.**
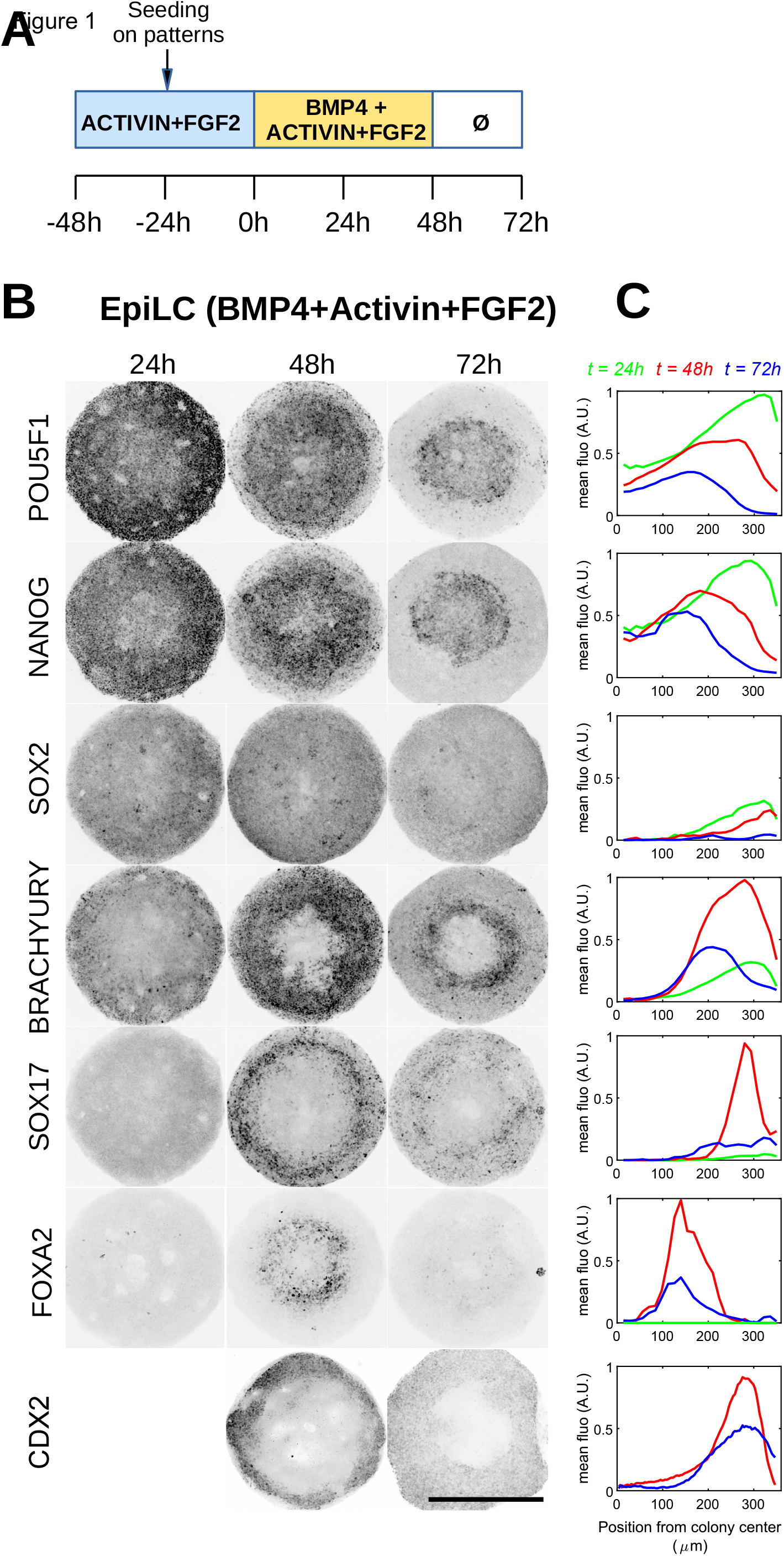
EpiLC colonies differentiation patterns by BMP4 stimulation. (A) Stimulation protocol to produce and differentiate EpiLC colonies on micropatterns. Ø indicates that no growth factors were added to the culture medium. (B) Maximum Intensity Projections of immunostaining of EpiLC colonies (700um) 1, 2, or 3 days after the beginning of BMP4 stimulation. For clarity, contrast in inverted. Each stain has been reproduced at least twice with similar results (C) Quantification of average fluorescence along the radii of colonies for t=24,48,72hr. POU5F1 (n = 18,24,24), NANOG (n=3,3,3), SOX2 (n=3,3,3), BRACHYURY (n=15,18,18), SOX17(n=3,6,6), FOXA2 (n=3,3,3) CDX2(3,3). For each marker, curves are normalized by the maximum of the set, except for SOX2 which is normalized by expression at t=0 (see Fig S1) CDX2 was not characterized at t=24h. scale bar = 500μm.

Pluripotency in the post-implantation embryo is known to track the expression of POU5F1 (OCT4), which becomes restricted to the posterior epiblast (Osorno et al., 2012). NANOG is re-expressed in the posterior epiblast while SOX2 expression, initially present in the entire epiblast, is progressively restricted to progenitors of anterior and neural fates after the onset of gastrulation (Avilion et al., 2003; Hart et al., 2004). We found that 24h after BMP4 addition EpiLC colonies expressed these three factors homogeneously (Fig. 1B). IFs at 48 and 72h showed that the expression of POU5F1 and NANOG then declined gradually and became restricted to the centre of the colonies, when that of SOX2 weakened and disappeared. These dynamics suggested the persistence of posterior epiblast cells in the centre of the colonies.

The pan-mesodermal marker BRACHYURY (BRA, also known as T), which starts to be expressed in the posterior epiblast at E6.0-E6.25 shortly before PS formation (Perea-Gomez et al., 2004; Rivera-Pérez and Magnuson, 2005), was detected in the colonies 24h after induction, in a broad outer ring of cells (Fig. 1B). This expression strengthened after 48h and moved gradually inward to reach a more central position at 72h. The expression of SOX17 was detected in a thin ring of cells within the BRA expression domain at 48h. These factors are both expressed in the embryo in extra-embryonic mesoderm cells contributing to the allantois, which emerge from the posterior PS, and in axial mesoderm cells emerging from the anterior PS(Burtscher and Lickert, 2009; Burtscher et al., 2012; Wilkinson et al., 1990). To find out which of these two possibilities fit with the pattern we obtained we looked at CDX2, which is co-expressed with BRA and SOX17 in some posterior mesoderm derivatives, and at FOXA2, which is co-expressed with BRA in posterior epiblast cells and axial mesodermal cells, and with SOX17 in definitive endoderm cells (DE) (Burtscher and Lickert, 2009; Burtscher et al., 2012). CDX2 was detected on the periphery of the colony at 48 and 72h. In contrast, FOXA2 expression was detected 48h after BMP4 addition in a ring of cells closer to the centre of the colonies, overlapping with NANOG-positive cells, but not with SOX17-positive cells. This indicated that FOXA2 expression in BMP4-stimulated colonies was associated with a posterior epiblast identity while SOX17 expression was associated with an extra-embryonic mesoderm identity. The fact that SOX17 and FOXA2 were not co-expressed and that FOXA2 expression was no longer detected 72h after BMP4 addition, strongly suggests that definitive endoderm and axial mesoderm do not form on BMP4-stimulated colonies.

These results thus showed that when exposed to BMP4 EpiLC colonies form a specific differentiation pattern, with a ring of mesoderm surrounding a core of epiblast, both biased towards posterior identities as embryological studies led us to expect (Kinder et al., 1999)

### 2. Ensuring the reproducibility of developmental trajectories

We found that the reproducibility of this differentiation pattern could be affected by a number of parameters. The first is the density of the adhesive micro-patterns on the glass slide. For high colony density, we noticed the existence of meta-patterns of differentiation, with colonies at the centre of each culture well displaying a staining pattern markedly different from that of colonies positioned at the edge of the well (Fig. S2A). This suggested that the diffusion of signalling molecules produced by the cells affected nearby colonies. This was confirmed when a reduction in the density of adhesive micropatterns resulted in all colonies adopting the same differentiation pattern regardless of their position in each well (Fig. S2B).

The second parameter affecting reproducibility is cell density. While differentiation patterns were highly reproducible within an experiment, occasional variations were associated with non-homogeneous cell seeding within a well or between wells. Likewise, variations between separate experiments were sometimes traced back to differences in the seeding density of the EpiLCs.

To push further our analysis of the reproducibility of the experiment required another approach than IF, where the number of markers that can be monitored at once is limited. We used data from transcriptome studies of gastrula stage embryos to select about 20 marker genes suitable to track the formation and patterning of the PS in BMP4-stimulated colonies (Peng et al., 2016; Pijuan-Sala et al., 2019) (Table S1). RT-PCRs were performed at 5 time points over a 72h culture on duplicates from a first experiment and on a second independent experiment performed another day. Genes were clustered in 3 groups according to the similarity of their expression dynamics in each set of colonies (Fig. S3A). In the first were epiblast markers – the pluripotency factors *Pou5f1* and *Sox2, Nodal* and its co-receptor *Cripto*, and the transcription factor *Otx2*, which is associated with epiblast maturation – whose expression was higher at the time of stimulation and tended to decline afterward. In the second were markers of the posterior epiblast and the primitive streak, such as *Nanog, Brachyury* and *Wnt3*, whose expression was activated 24h after stimulation. In the third were markers of the posterior PS and its mesoderm derivatives, such as *Cdx2*, *Gata4* and *Tbx4*, whose expression was activated at least 48h after stimulation. The resulting gene expression matrix confirmed the high reproducibility of the events taking place in the colonies after stimulation. Principal component analysis (PCA) of the data showed that while differences in the pace at which differentiation was taking place in the colonies could vary between independent experiments, their developmental trajectories nevertheless reached very similar endpoints (Fig. S3B), characterized by the expression of cluster 3 extraembryonic mesoderm markers. Another experiment showed that developmental trajectories were not significantly affected by the number of passages the cells had previously undergone (Fig. S3C, D).

We also tested the capacity of EpiSCs to generate differentiation patterns. First, we found that EpiSC colonies confined to adhesive micropatterns and stimulated with BMP4 failed to form a radially organized differentiation pattern and instead presented randomly distributed clusters of BRA-positive or SOX17-positive cells (Fig. S1D-F). This was reminiscent of the heterogeneity of EpiSCs in culture, which can be suppressed by treatment with the WNT secretion inhibitor IWP2 (Sugimoto et al., 2015). We then found that IWP2-treated EpiSCs have the capacity to form a radial differentiation pattern after BMP4 stimulation (Fig. S2B, C). RT-PCR analysis and PCA suggested that the developmental trajectories thus obtained were similar to those seen with EpiLC colonies (Fig. S1D). However, the need to treat EpiSCs with a pharmacological inhibitor to elicit a patterning response underlined how much more suitable EpiLCs were for our study, not to mention the availability of genetically modified ESCs.

These results thus confirmed that EpiLCs cultured on adhesive micro-patterns provide a robust system to study the molecular and cellular events underlying mouse epiblast and PS patterning. The characterisation of gene expression dynamics for a set of informative markers also proved a useful tool to track and to compare developmental trajectories, complementary to the visualization of differentiation patterns by immunofluorescence.

### 3. Sustained BMP exposure prevents the establishment of anterior cell identities

A recent study showed that EpiLC colonies like ours, when stimulated with a cocktail of BMP4, ACTIVIN, FGF2 and WNT3a, differentiated towards a mix of posterior cell identities quite similar to what we obtained with BMP4 alone (Morgani et al., 2018). This observation reinforced our interest to assess their response to ligands known to promote anterior cell identities in the embryo.

We thus examined the developmental trajectories of colonies stimulated with different combinations of the ACTIVIN (A), BMP4 (B), FGF2 (F) and WNT3a (W) ligands. We first compared the impact of no ligand (Ø), W, WAF and B on the colonies, and then we analysed the impact of BAF and BWAF to see how the presence of BMP4 affected the response to other ligand combinations. Samples were collected att=-48, -24, 0, 8, 24, 48 and 72h to track via RT-qPCR analysis the expression of 31 markers. They included the same genes as before, to which we added known targets of signalling pathways, such as *Axin2* (WNT/β-CATENIN target) and *Lefty2* (ACTIVIN/NODAL target), genes encoding secreted antagonists such as *Noggin*, *Chordin* and *Lefty2*, and several lineage markers, such as *Noto* and *Sox1b*

Genes were clustered according to the similarity of their expression dynamics (Fig. 2A). The higher number of markers and the establishment of anterior cell identities under certain conditions meant that the clustering was not identical to that described previously for BMP4-stimulated colonies. To summarize, cluster 1 markers track the disappearance of the epiblast identity, cluster 2 markers track the emergence of an anterior and neural identity and cluster 4 markers the emergence of extra-embryonic derivatives of the posterior PS. In contrast, cluster 3 markers can track the PS and its anterior (*Foxa2, Noto, Chd*) and posterior (*Pax3, Nog, Tbx6*) embryonic mesoderm derivatives. Our analysis made clear that, unlike BMP4, the W and WAF treatments promote the formation of anterior epiblast and anterior PS derivatives (cluster 2 and parts of cluster 3). The presence of BMP4 in the BAF and BWAF combinations however largely prevented the establishment of these identities, and thus appeared to block the influence of WNT and ACTIVIN/NODAL signalling on cell fate specification while promoting posterior fates (parts of cluster 3 and cluster 4). Interestingly, this does not appear to involve blocking these signalling pathways right away because the expression of *Wnt3* and *Nodal* as well as that of their respective feedback inhibitors, *Axin2* and *Lefty2*, was more strongly induced in the presence of BMP4.

**Figure 2.**
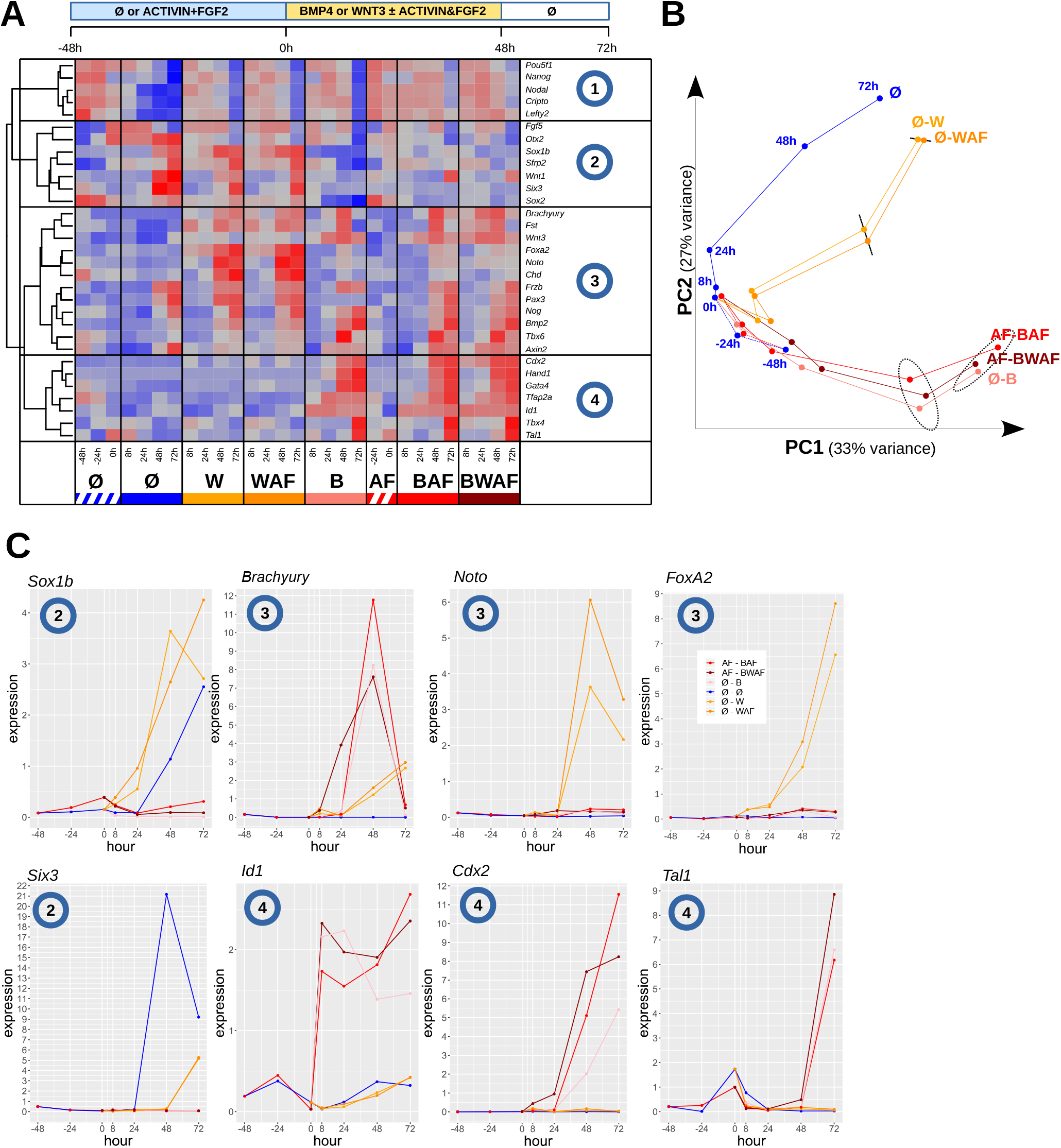
Comparison of temporal dynamics of EpiLC colonies differentiation depending on provided morphogens. (A) qRT-PCR expression matrix obtained for 2 different types of EpiLC differentiations (with or without ACTIVIN&FGF2 at -48/24/0h) and multiple combinations of stimulations at 8/24/48/72h (Ø=no growth factors; B=BMP4 50ng/ml, W=WNT3A, 200ng/ml; A=ACTIVIN, 20ng/ml; F=FGF2, 12ng/ml). Genes were clustered according to the similarity of their expression patterns in all conditions and 4 groups were defined based on the clustering. (B) Temporal trajectory of the different combinations of stimulation along the first 2 principal components. Dotted ellipses represent the 95% confidence area of the group formed by the BMP stimulated samples. (C) Individual temporal expression profiles of example genes belonging to clusters 2, 3 and 4.

To visualize the developmental trajectories, we projected the gene expression data in the plane formed by the first two principal components of the data set, capturing together about 60% of the variance (Fig. 2B). This analysis confirmed that all BMP4-stimulated samples followed the same developmental path. Similar results were obtained in a separate experiment (Fig. S3C, D). Given that the expression of *Wnt3* was induced within 8h of BMP4 stimulation, that *Nodal* expression was increased 24h after stimulation and that the addition of significant amounts of the WNT3a, ACTIVIN and FGF2 ligands failed to alter the developmental trajectories of BMP4-stimulated colonies, this strongly suggests that WNT3 and NODAL can only exert an influence on PS cell fate in the absence of BMP4.

The PCA also revealed that the addition of AF did not alter the developmental trajectory of WNT3a-induced colonies, even though it had the potential to increase the activity of the ACTIVIN/NODAL and FGF signalling pathways beyond what endogenous ligands normally achieve. Since we had confirmed a previous report that ACTIVIN was dispensable to obtain the conversion of mESCs into EpiLCs (Buecker et al., 2014), presumably because of endogenous NODAL production, some of the EpiLCs used in these experiments were obtained without it. In the absence of WNT or BMP stimulation, such as in the Ø condition, *Nodal* expression was not sustained in EpiLCs, confirming the essential requirement for these signalling activities upstream of *Nodal*, and the colonies differentiated towards anterior ectodermal and neural identities. However, we noticed in these colonies a complete absence of PS and PS derivative markers, which appeared to contrast with the presence of posterior mesoderm markers in some *Nodal^-/-^* embryos (Conlon et al., 1994).

These results are thus consistent with the embryological studies that supported the implication of BMP4, WNT3 and NODAL in the formation of the PS and their respective roles in the specification of posterior (BMP4) and anterior (WNT3 and NODAL) cell identities. They however identify situations where increases in the concentration of WNT, ACTIVIN and FGF ligands have no impact on the composition of the cell identities obtained at the end of the treatment, observations possibly at odds with the morphogen gradients hypothesis. They also show that sustained exposure to BMP4 prevents the establishment of anterior cell identities, thus defining BMP4 in the embryo not just as part of a positive feedback loop promoting the expression of *Nodal*, but also as part of a negative feedback loop locally blocking the impact of WNT and ACTIVIN/NODAL signalling on the expression of markers of anterior cell identities.

### 4. 2D mouse gastruloids recapitulate *Nodal* regulation in the epiblast

Two observations emphasized the importance of endogenous NODAL and led us to investigate the impact of its depletion on the differentiation of EpiLC colonies in greater details. The first was that the addition of ACTIVIN+FGF2 had no or little effect on the developmental trajectories of WNT3a or BMP4-stimulated colonies. The other was that non-stimulated colonies, which ceased to express *Nodal* shortly after their conversion, showed no evidence of forming any mesoderm (Fig. 2A).

First, using a Nodal-YFP reporter line - where one copy of the gene is mutated to express YFP instead of the ligand (Fig. 3A;(Papanayotou et al., 2014)) - we quantified via time-lapse imaging the spatio-temporal dynamics of *Nodal* expression in differentiating colonies. Stimulation with BMP4 or WNT3a both resulted within a few hours in a strong and homogeneous induction of *Nodal* expression in the entire colony (Fig. 3B). This expression peaked at t=24h and subsequently decreased, but not at the same speed everywhere so that it disappeared faster in the centre of the colony than on its periphery (Fig. 3C). In both BMP4 and WNT3a-stimulated colonies the ring of cells where *Nodal* expression persisted the longest was found to be part of the expression domain of the pan-mesodermal marker BRA (Fig. 1B; Fig. 3BC), as the colocalization of both markers in gastrula stage embryos led us to expect.

**Figure 3.**
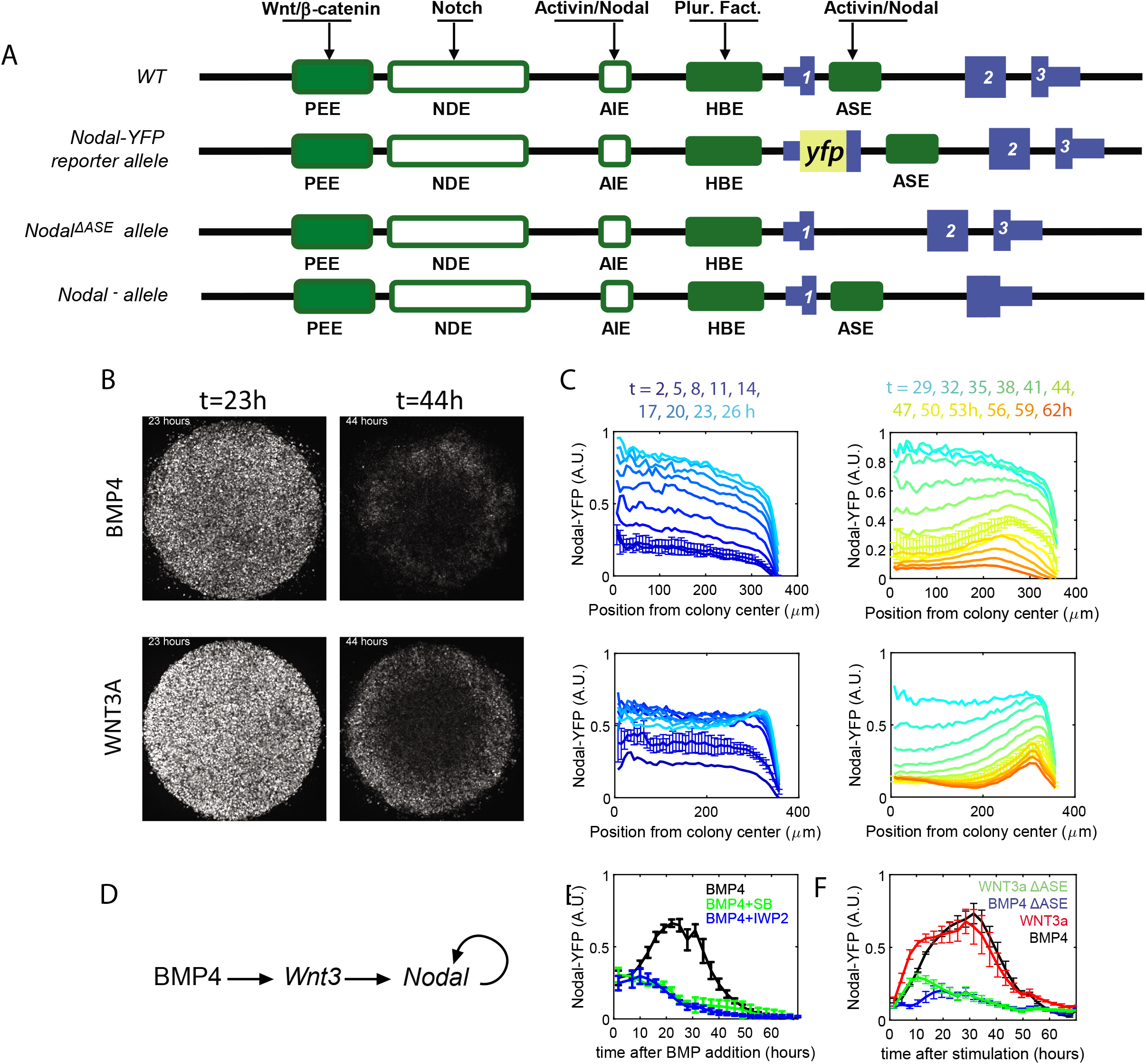
spatiotemporal profile of *Nodal* expression during EpiLC colonies differentiation. (A) Schematic view of the wt *Nodal* locus and of the modifications used in this study. The 3 exons are depicted in blue and regulatory elements as green boxes. Filled boxes represent regulatory elements that are active in the mouse embryo at epiblast and gastrulation stages. In one of the 2 alleles of the Nodal-YFP cell line, one of the 2 *Nodal* alleles has a yfp gene inserted in place of exon 1. In the *Nodal^ΔASE/ΔASE^* cell line, the ASE enhancer is deleted in both alleles. In the *Nodal^-/-^* cell line, parts of exon 2 and 3 are deleted, on both alleles. “Nodal-YFP ΔASE” line has ASE enhancer deleted on both alleles. (B) Expression of *Nodal* revealed by the Nodal-YFP reporter allele is homogenous in the colonies 24h after BMP or WNT stimulation and restricted in a ring in the periphery of the colonies at t=44h. (C) Evolution of the radial profile of Nodal-YFP expression as a function of time for BMP (top) or WNT (bottom) stimulation. profiles are averaged over n=4 colonies. For readibility, profiles of the first (left) and second (right) days are presented on two separate graphs and error bars representing standard deviation are only depicted for t=5h and 44h. (D) Simplified model of induction of *Nodal* on the posterior side of the mouse gastrula deduced form embryological studies. (E) Expression profiles averaged over n=4 colonies following BMP4 stimulation alone and with the inhibitors of ACTIVIN/NODAL pathway SB431542 (SB) or WNT secretion IWP-2. Average profiles of n= 4 colonies for each conditions. (F) Comparison of the Nodal-YFP expression temporal profiles following BMP4 or WNT3a stimulation for the unmodified Nodal-YFP line and the one where the ASE regulatory element has been deleted on both alleles (ΔASE). Average profiles of n= 4 colonies for each conditions. Experiments of each panel has been reproduced at least twice with similar results.

*Nodal* expression in the post-implantation epiblast is known to depend on two positive feedback loops. One involves the signalling cascade where BMP4 activates *Wnt3* and WNT3 activates *Nodal*, the other involves the promotion of *Nodal* expression by its signalling pathway (Fig. 3D). We found that the inhibition of ACTIVIN/NODAL signalling with SB431542 or of WNT secretion with IWP2 prevented the induction of *Nodal* expression in BMP4-stimulated colonies and led to its disappearance, confirming that this expression is like in the embryo dependent on both of these signalling pathways (Fig. 3E). Together with our previous observation that *Wnt3* expression is most induced in BMP4-stimulated colonies (Fig. 2A), these results are consistent with our 2D gastruloids adequately replicating the parts played by BMP4 and WNT3 upstream of *Nodal* expression in the post-implantation embryo. The auto-amplification of *Nodal* expression is mediated by the ACTIVIN/NODAL signalling-dependent *Nodal* enhancer ASE (Fig. 3A; (Norris et al., 2002; Yamamoto et al., 2001)). It has been shown recently that *Nodal* expression in preimplantation epiblast cells is initially under the control of another *Nodal* enhancer, called HBE, but that during epiblast maturation ASE becomes the predominant *Nodal* enhancer (Papanayotou et al., 2014). This regulatory shift is recapitulated *in vitro* during the conversion of ESCs into EpiLCs. Interestingly, mouse embryos homozygous for a deletion of ASE show no gastrulation phenotype, despite a drastic reduction in *Nodal* expression (Norris et al., 2002). It is only when combined with a KO allele of *Nodal* that the ASE-deleted *Nodal* allele results in gastrulation defects (Norris et al., 2002), suggesting that the use of such an hypomorphic allele might allow us to link *Nodal* expression levels and cell fate specification.

We thus used genome editing to generate homozygous deletions of the ASE in both *wt* and Nodal-YFP ESC lines, in order to characterize its contribution to the dynamics of *Nodal* expression in differentiating colonies. ASE-deleted cells showed reduction in the peak of Nodal-YFP expression after BMP4 or WNT3a stimulation of about 75%, relative to undeleted cells (Fig. 3F). ASE-deleted cells nevertheless showed a small increase in Nodal-YFP expression after stimulation, presumably mediated by the WNT signalling-dependent PEE since WNT-stimulated cells responded faster than BMP-stimulated ones.

2D gastruloïds thus correctly recapitulate *in vitro* the regulation of *Nodal* expression as it has been characterized in the embryo. These results also identify *Nodal^ΔASE/ΔASE^* cells as a convenient model to investigate how a hypomorphic allele of *Nodal* affects patterning.

### 5. *Nodal* is required to form posterior mesoderm in BMP4-stimulated colonies

To study the contribution of *Nodal* to colony patterning, in addition to the *Nodal^ΔASE/ΔASE^* cells described above, we also generated *Nodal^-/-^* cells by deleting a region that went from within exon2 to the beginning of exon3, thus preventing the production of an active ligand but leaving enough of the transcript to detect its expression. We then compared the differentiation of both mutants cell lines and of their unmodified parental cell line in BMP4-stimulated colonies. The markers we tracked were the same as before, and the analysis of their expression separated them in four clusters (Fig. 4A). In the first were early epiblast markers – including pluripotency factors and NODAL pathway components - whose expression tended to be higher before stimulation. In the second were markers of the maturing epiblast – *Otx2, Fgf5* and *Sfrp2* – that normally reached their peak just after stimulation. In the third were PS and posterior PS derivatives markers – such as *Bra, Cdx2* and *Tbx6* – which were most expressed 48h and 72h after stimulation in colonies of *wt* cells. In the fourth cluster were markers associated with a variety of cell identities, from mature epiblast and non-neural ectoderm to PS and cardiac mesoderm. Colonies of *Nodal^-/-^* cells, like non-stimulated *wt* colonies above, failed to activate *Bra* expression and showed no sign of forming a PS and mesoderm derivatives, adopting instead what we identified in cluster 4 as a signature of non-neural ectoderm. Again, this appeared different from the phenotype of *Nodal^-/-^* embryos where up to 25% were found to form random patches of posterior mesoderm (Conlon et al., 1994), implying that *Nodal* expression is more strictly required to form posterior mesoderm than the study of such embryos suggested.

**Figure 4.**
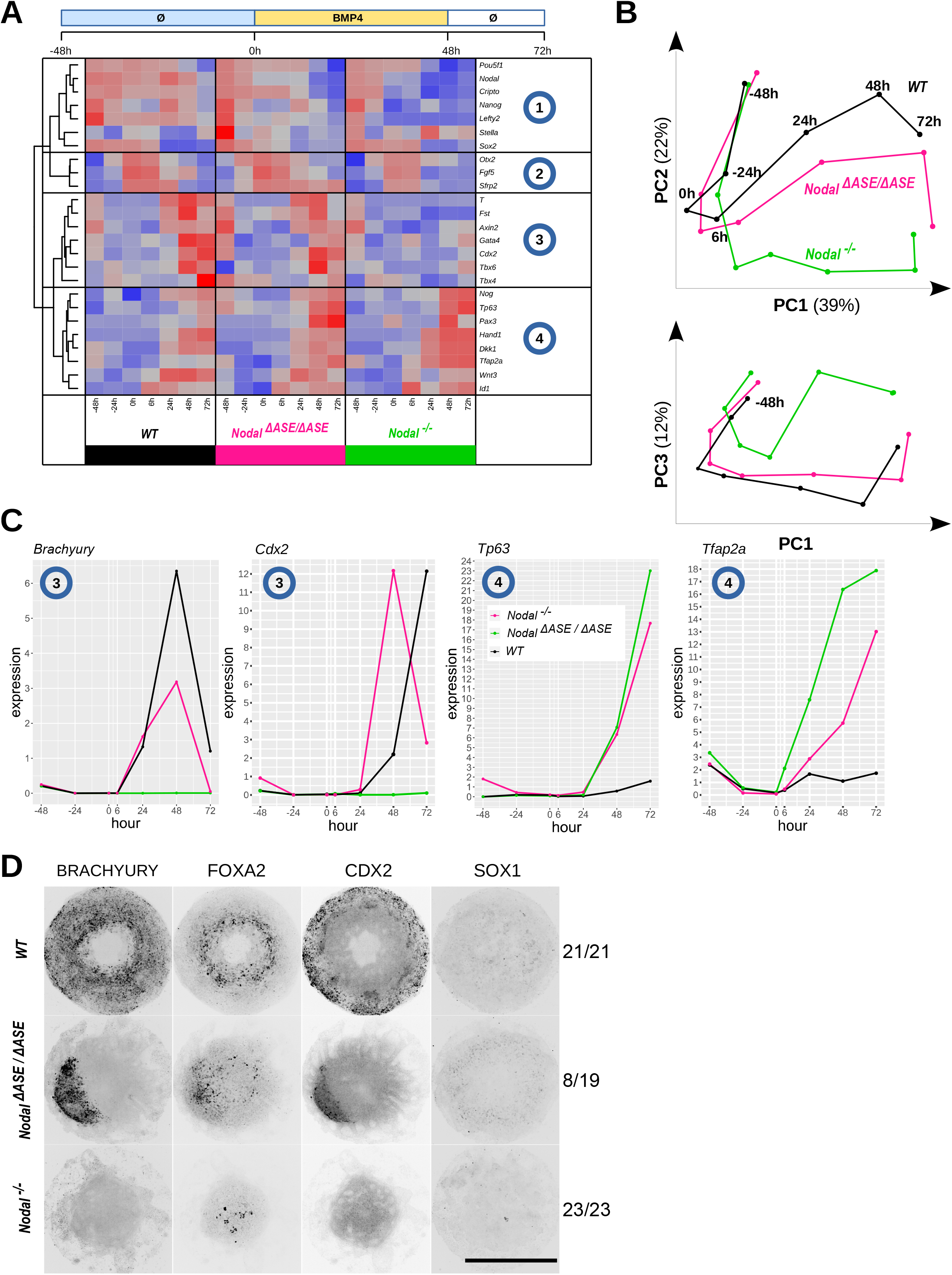
Comparison of wild-type and mutant colonies behavior after BMP4 stimulation. (A) qRT-PCR expression matrix obtained during EpiLC differentiation (without ACTIVIN&FGF2 at -48/24/0h) and after BMP4 stimulation at 6/24/48/72h for wild-type (WT) and mutant (*Nodal^ΔASE/ΔASE^* or *Nodal^-/-^*) cells. Genes were clustered according to the similarity of their expression patterns in all conditions and 4 groups were defined based on the clustering. (B) Temporal trajectory of the cell genotypes along the first 3 principal components, representing together 90% of the variance. (C) Individual expression profiles of example genes extracted from clusters 3 and 4 of the gene expression matrix presented in A. (D) Immunostainings of WT and mutant colonies after 48h of BMP4 stimulation. Over 2 separate experiments, 21/21 WT colonies displayed a full ring of BRACHYURY and no expression of BRACHYURY was detected in 23/23 *Nodal^-/-^* colonies. In *Nodal^ΔASE/ΔASE^* colonies, 4/19 colonies had a full ring of BRA, 6/19 a partial ring, 8/19 patches and 1/19 no expression. An example of the majority category (“patched”) is shown; see Figure S4 for examples images of BRACHYURY stains of each category. Scale bar 500μm.

Colonies of *Nodal^ΔASE/ΔASE^* cells also displayed a phenotype more severe than what embryonic phenotypes led us to expect. While they expressed *Bra* and other PS and posterior mesoderm markers, most peaked at 48h and declined afterward (Fig. 4C, S4A-C). Intriguingly, the expression of some cluster 4 genes (*Tp63, Pax3, Tfap2a, Id1*) appeared closer to what was seen in *Nodal^-/-^* colonies than in *wt* colonies (Fig. 4C). The developmental trajectories obtained by PCA showed indeed that *Nodal^-/-^* and *Nodal^ΔASE/ΔASE^* colonies reached endpoints that appeared closer than expected given how dissimilar the corresponding embryonic phenotypes were (Fig. 4B) (Conlon et al., 1994; Norris et al., 2002). Immunostaining of the different types of colonies for a small set of markers however revealed that while wt colonies showed the same radial differentiation pattern as before and *Nodal^-/-^* colonies showed no pattern at all, *Nodal^ΔASE/ΔASE^* colonies did not have a reproducible spatial organisation and most colonies expressed BRA, FOXA2 and CDX2 in irregular patches of variable size and number (Fig. 4D and S4D).

We then compared the differentiation of colonies from the same 3 cell lines after stimulation with WNT3a. Analysis of the gene expression dynamics of our panel of markers put them in 3 clusters (Fig. 5A). The first grouped together markers of early and maturing epiblast, from *Pou5f1* to *Fgf5*. The second contained anterior ectodermal and neural markers. The third contained PS and PS derivative markers, from *Bra* to *Cer1* (definitive endoderm). As before, WNT3a stimulation promoted in *wt* colonies the emergence of epiblast and PS derivatives of an anterior character. In *Nodal^-/-^* colonies anterior ectodermal and neural markers were more strongly induced, while the expression of most PS and PS derivatives markers remained at very low levels. A notable exception was *Cdx2*, which was transiently activated shortly after stimulation.

**Figure 5.**
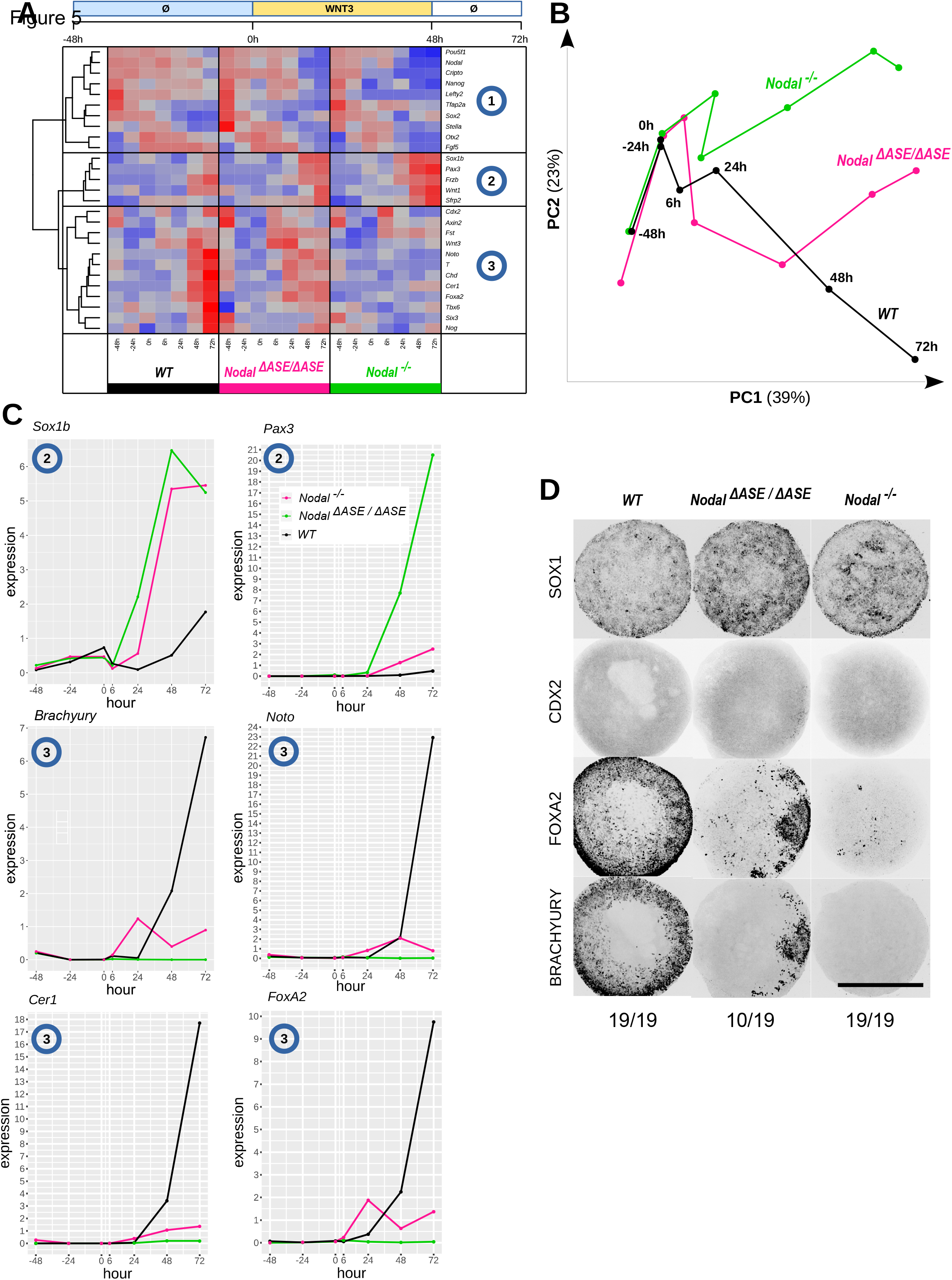
Comparison of wild-type and mutant colonies behavior after WNT3A stimulation. (A) qRT-PCR expression matrix obtained during EpiLC differentiation (without ACTIVIN&FGF2 at -48/24/0h) and after WNT3 stimulation at 6/24/48/72h for wild-type (WT) and mutant (*Nodal^ΔASE/ΔASE^* or *Nodal^-/-^*) cells. Genes were clustered according to the similarity of their expression patterns in all conditions and 3 groups were defined based on the clustering. (B) Temporal trajectory of the cell genotypes along the first 2 principal components, accounting for 60% of variance. (C) Individual expression profiles of example genes extracted from clusters 2 and 3 of the gene expression matrix presented in A. (D) Immunostainings of WT and mutant colonies after 48h of WNT3 stimulation. Over 2 separate experiments, 19/19 WT colonies displayed a full ring of BRACHYURY and no expression of BRACHYURY was detected in 19/19 *Nodal^-/-^* colonies. In *Nodal^ΔASE/ΔASE^* colonies, 2/19 colonies had a full ring of BRA, 5/19 a partial ring, 10/19 patches and 2/19 no expression. An example of the majority category (“patch”) is shown; see Figure S4 for examples images of BRACHYURY stains of each category. Scale bar 500μm.

*Nodal^ΔASE/ΔASE^* colonies showed again a stronger phenotype than expected, activating the expression of cluster 2 genes to levels similar to those of *Nodal^-/-^* colonies. Although they expressed higher levels of PS and anterior PS derivatives markers (such as *Bra, Noto, Cer1* and *Foxa2*) they did not reach the levels seen in *wt* colonies. In contrast, markers of anterior ectodermal and neural cells were upregulated, much like in *Nodal^-/-^* colonies. While the developmental trajectory of *Nodal^ΔASE/ΔASE^* colonies was initially closer to that of *wt* colonies, it became parallel to that of *Nodal^-/-^* colonies after 24h and its endpoint was quite different from that of either (Fig. 5B). Immunostaining showed in *Nodal^ΔASE/ΔASE^* colonies similar patches of BRA or FOXA2 expression as in BMP4-stimulated *Nodal^ΔASE/ΔASE^* colonies, with the same variability in size and expression levels (Fig. 5D; Fig. S4), only this time FOXA2 was clearly co-expressed with BRA and CDX2 was absent, as befit anterior PS derivatives. Interestingly, the drastic reduction in *Nodal* expression resulted in a reduction of the number of cells expressing anterior mesendoderm markers, but not in their replacement by cells expressing more posterior markers, such as *Tbx6*.

These results thus support a strict requirement for *Nodal* to endow EpiLCs with the capacity to form a PS and its derivatives. They show that the positive feedback loop mediated by ASE in the regulation of *Nodal* expression is critical to the robustness and reproducibility of the patterning. They also show that although *Nodal* expression is drastically reduced in *Nodal^ΔASE/ΔASE^* colonies, it is nevertheless sufficient to promote the expression of PS and posterior mesodermal markers in a fashion that is actually reminiscent of the variety of phenotypes characterized in *Nodal^-/-^* embryos (Ben-Haim et al., 2006; Brennan et al., 2001; Conlon et al., 1994), suggesting the presence in some of the ligands that partially compensate for the absence of endogenous NODAL.

### 6. ACTIVIN of extra-embryonic or decidual origin may partially rescue the phenotype of *Nodal^-/-^* embryos

Our results suggest that the presence of posterior mesoderm derivatives in some *Nodal^-/-^* embryos is due to the persistence of BMP4 ligands, and this interpretation is supported by reports of residual *Bmp4* expression in the ExE of E5.5 and E6.5 *Nodal^-/-^* embryos (Brennan et al., 2001; Mesnard, 2006). They also suggest that the epiblast of these embryos has been exposed to ACTIVIN/NODAL-like ligands. *Nodal* is expressed in the uterus at postimplantation stages but in cells at positions quite remote from the embryo (Park and Dufort, 2011). Furthermore, *Nodal^-/-^* embryos rapidly loose *Cripto* expression and therefore the capacity to respond to NODAL signalling (Brennan et al., 2001; Mesnard, 2006). In contrast, ACTIVIN subunits are expressed in the decidual cells that surround the conceptus at these stages and at low levels in extra-embryonic tissues (Albano et al., 1994; Pijuan-Sala et al., 2019), and ACTIVINs do not need CRIPTO to activate downstream signalling. RT-qPCR analyses confirmed that *Nodal^-/-^* colonies recapitulate the loss of *Cripto* expression known to take place in mutant embryos (Fig. 6A). We thus attempted the rescue of the patterning of BMP4-stimulated *Nodal^-/-^* colonies either with recombinant NODAL or with ACTIVIN. We found as expected that these colonies failed to respond to NODAL but did express BRA when treated with ACTIVIN (Fig. 6B). This result suggests that the presence of ACTIVIN of extra-embryonic or maternal origin in *Nodal^-/-^* embryos is a possible explanation for the apparent discrepancy between their patterning defects and those we characterized in *Nodal^-/-^* EpiLC colonies.

**Figure 6.**
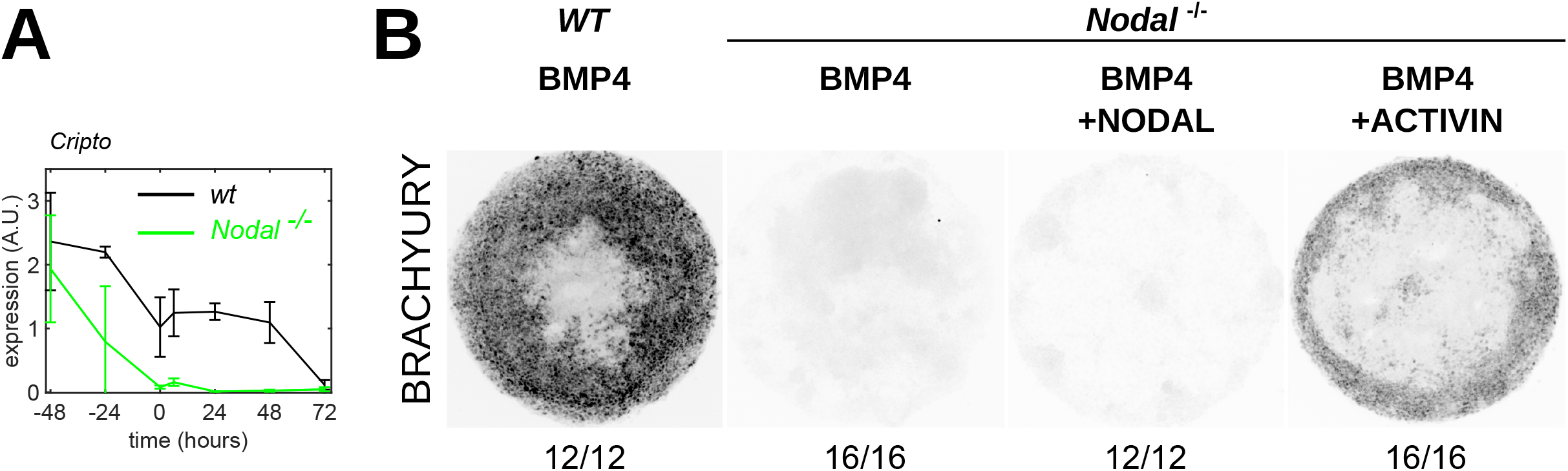
Rescue of differentiation patterns in Nodal KO mutants with NODAL or ACTIVIN. (A)BRACHYURY immunostainings of *wt* and *Nodal^-/-^* mutant colonies after 48h of stimulation with BMP4 (50ng/ml) alone or in combination with NODAL (200ng/ml) or ACTIVIN (20ng/ml). Over 2 separate experiments, no BRACHYURY expression was detected in colonies treated by BMP4 (n=12) and BMP4+NODAL (n=12), while BRACHYURY expression was rescued in colonies treated with BMP4+ACTIVIN (n=16). (B) Expression of *Cripto* during *wt* and *Nodal^-/-^* EpilC differentiation. BMP4 (50ng.ml) was added to the medium at t=0. Average profile over n=3 experiments, expression values normalized by expression at t=0, error bar: standard deviation.Plouhinec et al.

## Discussion

We have adapted to the use of mouse pluripotent stem cells an *in vitro* approach for the study of embryo patterning that was first developed using human cells (Warmflash et al., 2014). We identified conditions that ensure the reproducibility of the differentiation events taking place in micropatterned EpiLC colonies after their exposure to different morphogens. Although these conditions were slightly different from those recently described by another group, the differentiation patterns we obtained were similar to theirs (Morgani et al., 2018), further attesting of the robustness of the approach. We used immuno-fluorescence, RT-qPCR and PCA to characterize the differentiation patterns thus generated and to compare the developmental trajectories induced by distinct combinations of signalling molecules. The results of these experiments are broadly consistent with embryological data that support a role for BMP4 in promoting ectodermal and posterior mesodermal fates and a role for NODAL and WNT3 in promoting anterior mesendodermal fates. However, they also show that the presence of BMP4 prevents the emergence of anterior cell fates, regardless of the presence of ACTIVIN/NODAL and WNT ligands and their ongoing signalling. Furthermore, they show that NODAL is strictly required to form a PS and all its derivatives and that a drastic reduction in its expression level does not result in a shift to the formation of more posterior PS derivatives.

### BMP has a dominant effect on colony fate

BMP4-stimulated colonies formed a ring of posterior mesoderm cells surrounding a core of non-neural ectoderm and this pattern was consistent with the defects described in *Bmp4^-/-^* and *Bmpr1a^-/-^* embryos where the formation of these cell identities is defective (Di-Gregorio et al., 2007; Winnier et al., 1995). The addition of WNT3a and ACTIVIN to BMP4 in the differentiation medium did not change the colonies’ developmental trajectories. Since BMP4 induces both *Wnt3* and *Nodal* expression in the colonies, as in the posterior epiblast, this suggests that their respective signalling activities are tightly regulated and not affected by the addition of exogenous ligands. However, we found that in the absence of BMP4, ACTIVIN and WNT3a do promote the formation of anterior derivatives of the PS and of neural fates. This strongly suggests that BMP4, in addition to promoting posterior cell fates, actively suppresses anterior ones. There are several examples of BMP and NODAL signalling antagonizing each other during embryogenesis (Furtado et al., 2008; Pereira et al., 2012; Yamamoto et al., 2009). These situations emerge when a component common to both signalling pathways, such as SMAD4, becomes limiting and competition leads to reduced output. A similar situation is suspected in the PS but it is not clear that it fits what takes place in our colonies. On one hand, *Id1* expression following BMP4 stimulation was much stronger in *Nodal^-/-^* EpiLC colonies than in *wt* ones. On the other, the expression of *Nodal* and of its target gene and feedback antagonist, *Lefty2*, appeared unchanged or even slightly higher in *wt* colonies exposed to BMP4. The WNT signalling pathway appeared likewise unaffected. This suggests a more selective repression, presumably at the level of the chromatin or gene regulatory networks, of the WNT and NODAL signalling targets that are critical to the emergence of the anterior PS and its derivatives. Alternatively, the changes in gene expression induced in BMP-responsive cells may provide a context in which antagonism to NODAL and WNT signalling becomes gradually more effective.

In the absence of BMP4, WNT with or without ACTIVIN+FGF2 promotes in EpiLC colonies the formation of anterior PS derivatives, a result consistent with their emergence in the embryo from a position in the PS that is the further away from BMP influence and where BMP antagonists are secreted. The failure of colonies that are exposed to BMP4 to form anterior PS derivatives may well result from its continued presence. In the embryo cells ingress through the PS and then move away from it, and their exposure to some of its signals is only transient. It would therefore be interesting to assess how shorter exposures to BMP4 would affect the developmental trajectories of the colonies and the mix of cell identities they give rise to.

### NODAL drives a binary fate choice between mesendodermal and ectodermal identities

*Nodal* was found to be required for the formation of the PS (Conlon et al., 1991, 1994; Iannaccone et al., 1992; Zhou et al., 1993), but it was subsequently shown that *Nodal^-/-^* embryos also lacked *Wnt3* expression while that of *Bmp4* was drastically downregulated (Brennan et al., 2001). Both of these factors being also required to form a PS (Liu et al., 1999; Winnier et al., 1995), this raised the question of the respective contributions of each ligand to the process. The detection in some *Nodal^-/-^* embryos of cells of a posterior mesoderm character, expressing *Bra* or *Cdx4*, was originally ground for the claim that NODAL was required to form the PS but not these cell populations (Conlon et al., 1994). The fact that *Nodal^-/-^* EpiLC colonies when stimulated with BMP4 or WNT3a formed ectodermal derivatives of a posterior or an anterior character, respectively, but failed to express markers of the PS and its derivatives, highlights a strict requirement for NODAL for both PS formation and the specification of a mesendodermal identity. This is consistent with other embryological studies that found no evidence of the expression of PS or nascent mesoderm markers in *Nodal^-/-^* embryos (Ben-Haim et al., 2006; Brennan et al., 2001). The fact that colonies deprived of NODAL form anterior ectodermal and neural cell identities regardless of the presence of WNT ligands is also consistent with the presence of the same cell identities in *Nodal^-/-^* embryos (Camus et al., 2006). These results position NODAL as the determining factor in a binary choice between ectodermal and mesendodermal identities. They are consistent with NODAL acting upstream of the Tbx factors EOMES and BRACHYURY, which have been shown recently to govern the same binary choice via their impact on chromatin state (Tosic et al., 2019).

### *Nodal* mutant phenotypes may be partially rescued by ligands of extra-embryonic or maternal origin

While embryos homozygous for a deletion of the *Nodal* auto-regulatory enhancer ASE showed no gastrulation phenotype (Norris et al., 2002), EpiLC colonies carrying a similar deletion were unable to form radial differentiation patterns after stimulation with BMP4 or WNT3a. The elimination of ASE resulted in a 50 to 80% reduction in *Nodal* expression after stimulation. Feed-forward mechanisms are known to ensure robustness and reproducibility to the processes in which they are involved. Consistent with this view, the random patches of mesoderm that formed on *Nodal^ΔASE/ΔASE^* colonies attest of a disrupted patterning mechanism. The fact that nothing similar has been described in *Nodal^ΔASE/ΔASE^* embryos suggests that other ligands are compensating for the depletion of endogenous NODAL in these embryos, something the milder discrepancy between the *Nodal^-/-^* embryo phenotype and the patterning defects of *Nodal^-/-^* EpiLC colonies also hinted at. One obvious candidate is the ACTIVIN produced by extra-embryonic and decidual cells at the relevant post-implantation stages (Albano et al., 1994). The fact that the addition of ACTIVIN, but not of NODAL, could partially rescue the patterning of *Nodal^ΔASE/ΔASE^* colonies support this possibility and is consistent with the lack of expression of the NODAL co-receptor *Cripto* in *Nodal^-/-^* embryos and in *Nodal^-/-^* colonies.

### Possible inconsistencies with the A/P Nodal activity gradient hypothesis

A strict interpretation of the hypothesis stating that an A/P gradient of NODAL signalling activity patterns the P/S would lead us to expect that a lower expression of *Nodal* would result in the formation of more posterior mesodermal cell identities. This is what has been observed in human 2D gastruloids, where combined ACTIVIN and WNT stimulation promoted more anterior fates than WNT alone and inhibiting the ACTIVIN/NODAL pathway in that context lead to the expression of more posterior mesoderm markers, such as TBX6 (Martyn et al., 2019), although this last condition as also be shown to promote neural crest identity (Funa et al., 2015). In EpiLC colonies, by contrast, WNT3a stimulation alone was sufficient to induce anterior markers (*Noto, Cer1, Foxa2*) and addition of ACTIVIN only moderately increased their expression. Conversely, reducing *Nodal* expression by deleting its ASE enhancer did not result in a shift toward more posterior PS identity, but instead it promoted ectodermal markers. Together with the observation that patches of endoderm do get formed on *Nodal^ΔASE/ΔASE^* colonies treated with WNT3a, despite *Nodal* expression in these colonies remaining well below its normal levels, these observation are challenging the view, supported by many embryological studies, that the specification of the definitive endoderm calls for the highest levels of NODAL signalling (Dunn et al., 2004; Robertson, 2014). Our observations could result from the fact that NODAL signaling was reduced too much in *Nodal^ΔASE/ΔASE^* colonies to reveal gradation or that the graded effect becomes effective only in late primitive streak, a regime that EpiLC may fail to probe as they are close to the E5.5 epiblast when they receive gastrulation inducing signals in our experiments.

Nevertheless, there is clear evidence that NODAL signals are not conveyed in the same manner at all levels of the PS, implying that the efficiency of the signalling may vary. SMAD4, a key transducer of both ACTIVIN/NODAL signalling and BMP signalling is thus required in the epiblast to form derivatives of the anterior PS but not of mesoderm populations originating at more posterior levels (Chu et al., 2004). The formation of endoderm in *Nodal^ΔASE/ΔASE^* colonies suggests that the choice of how NODAL signalling is transduced and interpreted is actually not strictly dependent on the level of *Nodal* expression. In contrast, it could be affected by the presence of BMP4. Overall, our results highlight the need to characterise the spatio-temporal activity of the ACTIVIN/NODAL signalling pathway in the mouse gastrula, in order to clarify its role in the patterning of the PS.

## Experimental procedures

### Cell culture and cell lines

For *wt* mESCs, we used the HM1 line (Selfridge et al., 1992). Nodal::YFP line was established in a CK35 background (Papanayotou et al., 2014). mESC were cultured on 0.15% gelatin coated tissue culture grade plates in N2B27+2i+LIF medium (Silva et al., 2008). *Nodal^-/-^* and *Nodal^ΔASE/ΔASE^* line were established in HM1 background using CRISPR gene editing. For the *Nodal^-/-^* line, a 1196bp deletion spanning parts of exons 2 and 3 includes most of the propeptide. The Nodal^ΔASE/ΔASE^ line, reproduces the 600bp ASE deletion described in the literature (Norris et al., 2002).

### EpiLCs differentiation on micropatterned adhesive substrates

ESC cultures in N2B27+2i+LIF were harvested by trypsinization and seeded cells were first cultured for 24h on plates coated with fibronectin (15ug/ml for 30 min) in N2B27 supplemented with 1% KSR and optionally with 20ng/ml ACTIVIN and 12ng/ml FGF2. After 24h, cells were dissociated using trypsin and seeded on micropatterned substrates, produced by micro-contact printing fibronectin on PDMS-coated glass cover slip, at a density of 8000 cells per mm^2^. After 1h, cells that did not attach were removed by gentle flushing and allowed to form colonies for an additional 24h before being stimulated by adding different combinations of factors to the medium (BMP4:50ng/ml, WNT3a:200ng/ml, ACTIVIN: 20ng/ml and FGF2:12ng/ml). The medium was renewed every 24h during the course of the experiment.

### Quantitative analysis of gene expression

For each time point and conditions, cells from one well, containing between 7 and 10 colonies, were dissociated in 1M guanidium for further quantification of gene expression using standard qRT-PCR methods, see supplement for technical details. For each condition, duplicates samples were analyzed. Each experiment has been reproduced independently at least twice.

A custom R script was used to compute the following steps. In order to remove genes with low expression from later analysis, we removed genes which had a difference of at least 10 between their cycle quantification (Cq) value and the one of GAPDH in all samples considered. Then we computed for each sample and gene a relative gene expression value with respect to the mean expression of the gene in the experiment (obtained by pooling all samples belonging to the same experiment) and normalized by the expression of the GAPDH gene in the sample. This value was log2-transformed, then centered and reduced with respect to the expression value of the gene in all samples considered in order to compute and display the expression matrix and the principal component analysis of this matrix.

See SI for a detailed description of the procedures, a complete list of reagents, primers and antibodies used.

## supplementary materials

**Figure S1 (related to figure 1):**
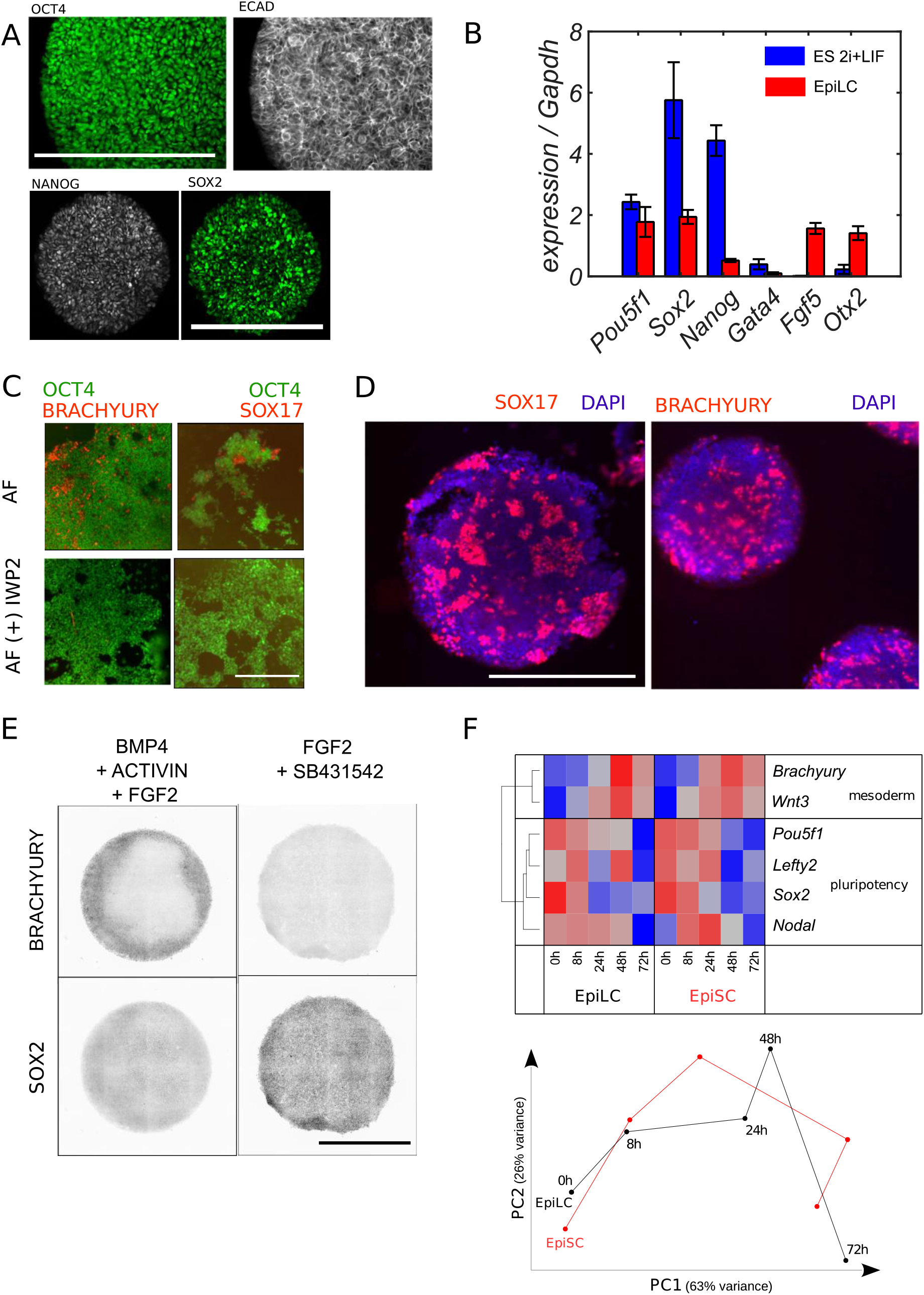
Characterization of EpiLC and differentiation of EpiSC on micropatterns. A - EpiLC characterization 48h after replacement of 2i+LIF by in ACTIVIN+FGF homogenous stain for OCT4, ECAD, NANOG and SOX2. Representative image of n=7 pictures. B – qRT-PCR characterization of the expression of markers characteristic of naïve and formative pluripotency, for mESC maintained in N2B27+2i+LIF medium and for micropatterned EpiLC colonies 48h after initiation of their conversion in either N2B27+ACTIVIN+FGF+KSR (EpiLC). The EpiLC conversion is characterized by down-regulation of core pluripotency markers *Nanog* and *Sox2* but not *Pou5f1*, upregulation of epiblast markers *Fgf5* and *Otx2*, while primitive endoderm markers such as Gata4 remains low, as previously reported in (Hayashi et al., 2011) error bars: standart deviation of n=2 separate experiments C - EpiSC maintained in growth medium containing ACTIVIN and FGF (AF) are heterogeneous and display cluster of cells expressing anterior primitive streak markers BRA and SOX17. (top row) (patches of variable size were observed in 10/10 randomly selected fields). This heterogeneity is not observed after 3 passages if an inhibitor of WNT secretion (IWP2) is added to the growth medium (bottom row) (no BRA or SOX17 patches observed in 10/10 randomly selected fields) D - EpiSC maintained in ACTIVN +FGF medium display patchy differentiation after 48h of BMP4 differentiation on micro-patterned substrates (n=60 colonies, 2 replicates) E - EpiSC maintained in AF+IWP2 for 3 passages before being seeded on micropatterned substrates and differentiated for 48h in 50ng/ml BMP4 stimulation display a radial pattern of differentiation (left column). Right column: the ring of BRACHYURY is lost upon ACTIVIN/NODAL pathway inhibition with the small molecule inhibitor SB431542 (SB). Representative pictures of n=4 colonies for each condition. F - Comparison of gene expression temporal dynamics between EpiSC and EpiLC during BMP4 induced differentiation on micro-patterns. (Gene expression matrix and temporal trajectory along the first 2 principal components). Scale bars: 500μm

**Figure S2 (related to Figure 1):**
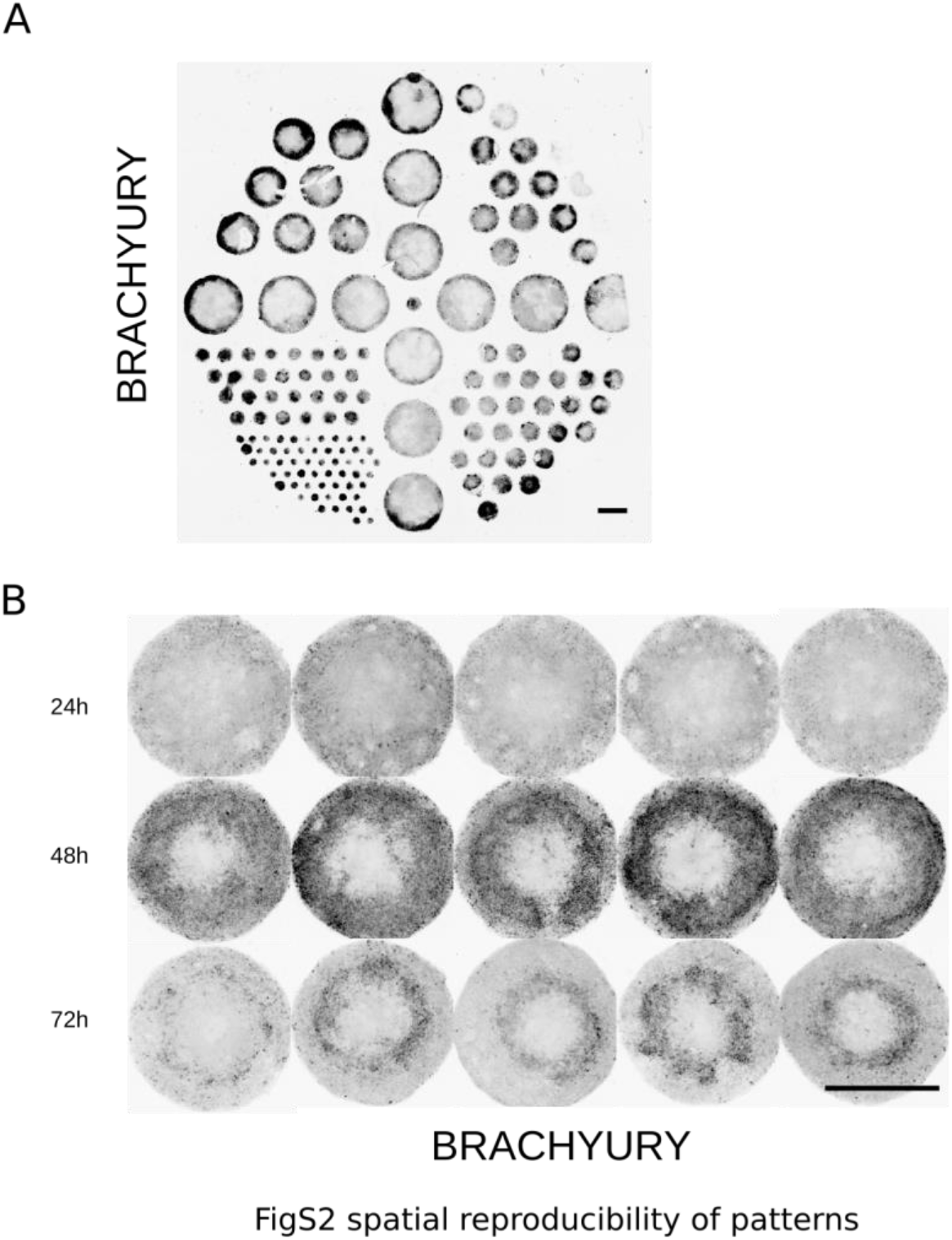
Quantification of the reproducibility of spatial organization of colonies under BMP4 induced differentiation. (A) A meta-pattern of differentiation is observed inside a well if colony density is too high and if original cell seeding is spatially homogeneous. See that BRACHYURY stain is stronger for colony on the edges of the well-cells were fixed and stained after 48h of BMP4 differentiation (B) BRACHYURY immunostaining of multiple EpiLC colonies cultured with ACTIVIN and FGF2 and stimulated with BMP4 for 48h. For low colony density (each colony is separated from its closest neighbor by two colony diameters) and homogenous seeding, the observed pattern is highly reproducible inside a well. Scale bars: 500μm

**Figure S3 (related to Figure 2):**
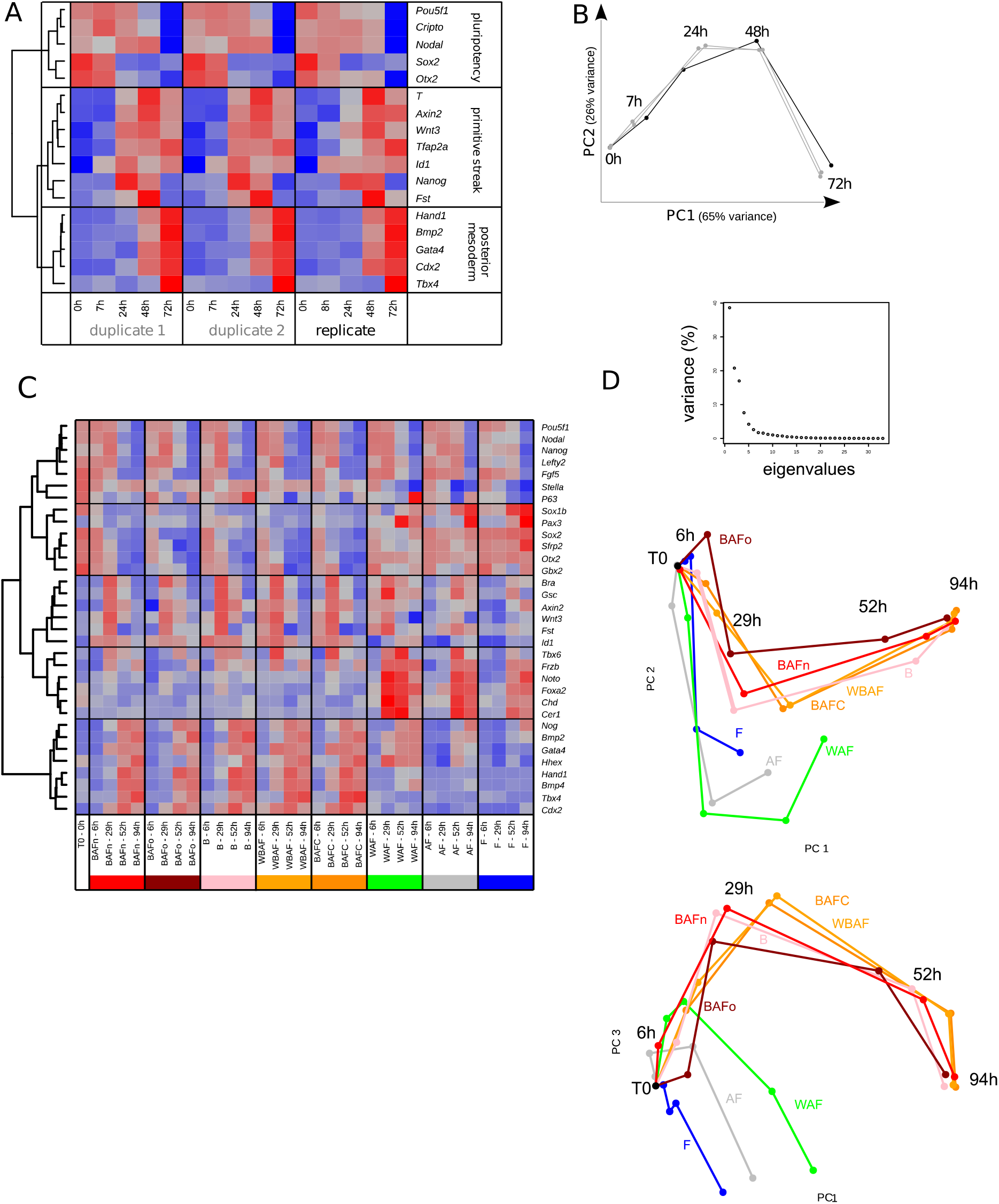
characterization of reproducibility of developmental trajectories. (A) Gene expression matrix of multiple replicates obtained by qPCR 0/7/24/48/72h after BMP stimulation. Duplicates 1&2 belong to the same experiment. The replicate belongs to and independent experiment. Genes were clustered according to the similarity of their expression patterns in all replicates and 3 groups were defined based on the clustering. (B,) Temporal trajectory of the replicates along the first 2 principal components. (C,D) replicate experiment of figure 2. (C) Gene expression matrix obtained by qRT-PCR 0/6/29/52/94h after various stimulations: B: BMP4, A:ACTIVIN, F:FGF2, W: WNT3a. BAFn and BAFo are two replicates of the same protocol but with cells with not the same passage number (n: 4 passages in N2B27+2i+LIF, o:27 passages in N2B27+2i+LIF) all other conditions are done with early passage cells

**Figure S4 (related to figure 4):**
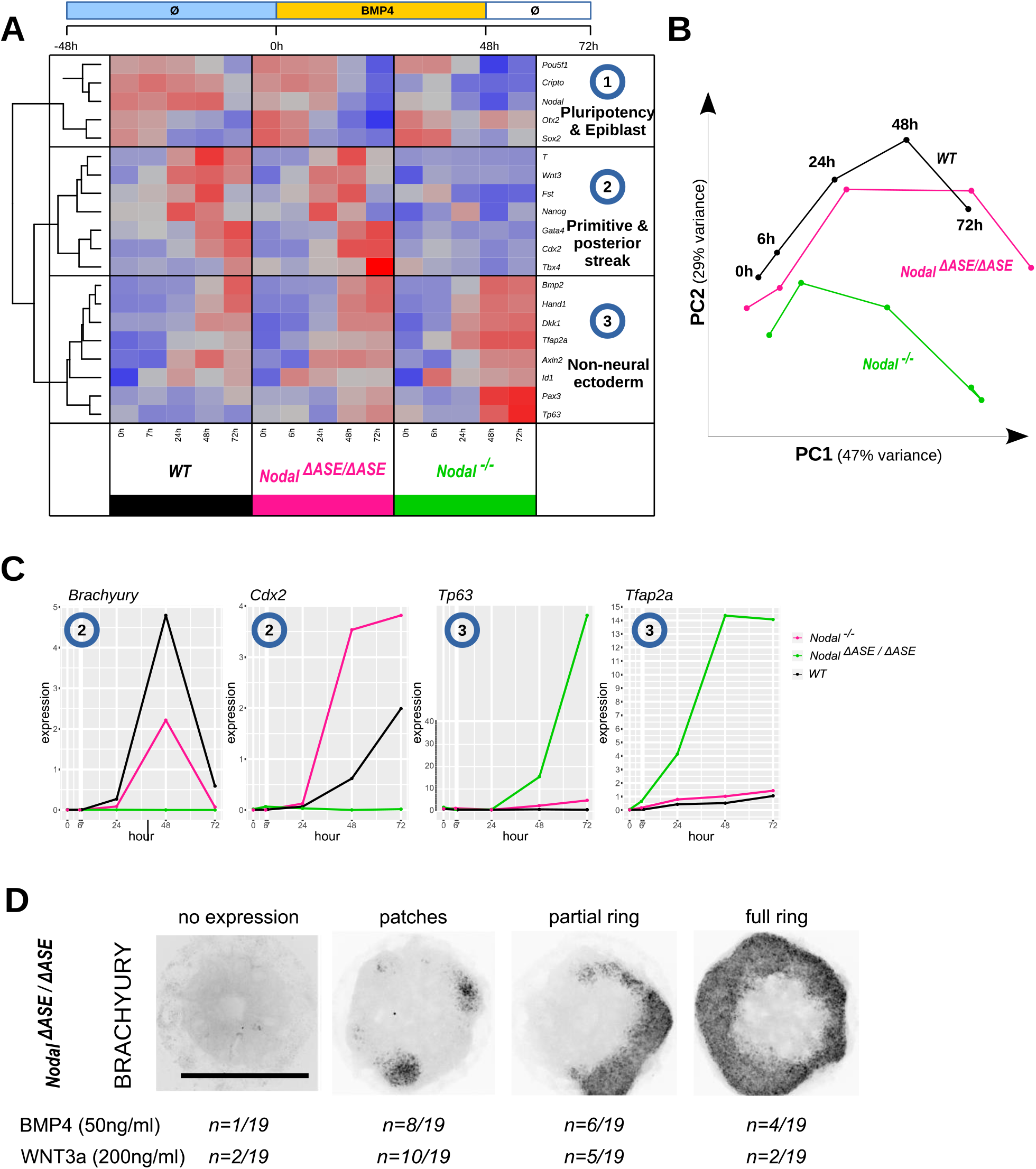
Experimental replicate of wild-type and mutant colonies after BMP4 stimulation. (A) qRT-PCR expression matrix obtained after BMP4 stimulation at 0/7/24/48/72h for wild-type (WT) and *Nodal* mutant cells; *Nodal^ΔASE/ΔASE^* (ASEKO) or *Nodal^-/-^* (Nodal KO). Genes were clustered according to the similarity of their expression patterns in all conditions and 3 groups were defined based on the clustering. (B) Temporal trajectory of the cell genotypes along the first 2 principal components. (C) Individual expression profiles of example genes belonging to clusters 2 and 3. (D) Immunostaining against BRACHYURY of BMP stimulated *Nodal^ΔASE/ΔASE^* (ASE KO) colonies. BRACHYURY expression is spatially heterogeneous with one or a few patches of positive cells per colony. Scale bars: 500μm

**Table S1 (related to figures 2,4&5): average single cell expression of markers used in this study in the mouse gastrula, arranged by tissue types**

These data were extracted from the publicly available dataset from (Pijuan-Sala et al., 2019)

**Cell types form the public data sets have been aggregated the following way:**

**“Caudal epiblast+”** = “Caudal epiblast”, “Caudal Mesoderm”
**“Nascent mesoderm+”** = “Nascent mesoderm”, “Mixed mesoderm”
**“Pharyngeal mesoderm+”** = “Pharyngeal mesoderm”, “Cardiomyocytes”
**“ExE mesoderm+”** = “ExE mesoderm”, “Allantois”, “Mesenchyme”
**“Haematoendothelial progenitors+”** = “Haematoendothelial progenitors”, “Endothelium”, “Blood progenitors 1/2”, “Erythroid1/2/3”
**“Def. endoderm+”** = “Def. endoderm”, “Notochord”, “Gut”
**“Neurectoderm+”** = “Caudal neurectoderm”,”Rostral neurectoderm”, “Forebrain/Midbrain/Hindbrain”, “Spinal cord”,”Neural crest”

## Extended Materials and Methods

### Cell culture

We used mESCs line HM1 (Selfridge et al., 1992) for *wt* cells and *Nodal^-/-^* and *Nodal^ΔASE/ΔASE^* mutants. Nodal-YFP line was established in a CK35 background (Papanayotou et al., 2014). mESC were cultured on 0.15% gelatin coated tissue culture grade plates in N2B27 medium (DMEM/F12 and Neurobasal media (1:1, Life Technologies) supplemented with 1× B27 1×N2 (Life Technologies), 2 mM L-glutamine (Life Technologies), 0.1 mM β-mercaptoethanol, penicillin and streptomycin (Life Technologies) LIF [1000U/ml cell guidance systems GFM200], PD0325901 [1μM, cell guidance systems, SM26] and CHIR99021 [3μM, cell guidance systems, SM13]. They were passaged every 2-3 days by dissociating them in 0.05% Trypsin [gibco, 25300-054] for 5 min, neutralizing trypsin with 15% serum supplemented DMEM, concentrating cells by spinning, and transferring to a new plate.

EpiSC cells (line FT129_1, a gift from Alice Jouneau) were cultured on serum coated plates in the following medium: IMDM/F12 50:50 [Invitrogen 31980-32 and 31765-027] supplemented with 0.5% BSA [life technologies15260-037], 1% lipid supplement [Invitrogen 11905-031], 450uM Monothioglycerol [Sigma M6145], 7μg/ml Insulin [Roche, 11376497001], 15μg/ml Transferin [Sigma T8158], 20ng/ml Activin [cell guidance systems, GFM29], and 12ng/ml FGF2 [cell guidance systems, GFM12],. They were passaged with 1mg/ml collagenase IV[life technologies, 17104-019] in DMEM/F12 [life technologies 21331] followed by colony fragmentation by pipetting.

### Generation of Nodal^-/-^ and Nodal^ΔASE/ΔASE^ cell lines

mESC mutant lines and were established using a CRISPR plasmid containing guide RNA and a Cas9-2A-OFP (Life Technologies A2117). The choice of gRNA was made using the CRISPR MIT tool (http://crispr.mit.edu/ of The Laboratory of Professor Zhang (Hsu, Lander and Zhang, 2014)) The targets selected in this study were chosen with a potential off-target score greater than 80/100 to minimize the risk of adverse effects. For the Nodal^-/-^ line, we used 5’ GTCGAGCAGAAAAGTGTTGG and 3’ ACCGGGTTCCTTCCACGTGC as gRNA pair. The 1196bp deletion includes part of the exon2 until the beginning of the exon3, so a whole intron and a piece of exons 2 and 3 is missing, including the major part of the region coding for Nodal propeptide. For the Nodal^ΔASE/ΔASE^ line, we used ACGATTTCTAAACTACAGAT and CGGCGGGCGGCGGGTCAGAC as gRNAs to reproduce the ASE deletion described in the literature (Norris et al., 2002). The cloning of gRNAs into the plasmid was carried out following the recommendations of the manufacturer. For transfections, 500,000 cells were seeded in a 35mm diameter petri dish in 1.5ml of ES-serum medium (DMEM+10% FBS+ LIF), 1 hour before the addition of transfecting agent. A mixture of 3μl of lipofectamine2000 and 1μg of each transfected plasmid was prepared in parallel in 1ml of DMEM. The mixture was left to incubate 30 minutes before being dropped on the cells. The medium was replaced 24 hours after the transfection. 48 hours after transfection, Fluorescent cells were FACS-sorted to select cell that received the plasmid. These are then seeded at low-density (1000 to 5000 cells per petri dish 10cm in diameter) in ES-serum medium. When the colonies are large enough, they were picked manually and further expanded in 96 wells plates. After passing, a fraction of each well was recovered in order to perform the genotyping of the cells. DNA extraction was achieved using Sigma RedExtract-N-Amp for tissue (Sigma XNAT-1K) kit, following manufacturer’s instructions

Genotyping was performed by PCR reaction, using PrimeSTAR GXL buffer, 1.2μl dNTP Mixture TAKARA, 0.2μl of each primer at 10 μM, 1μl of PrimeSTAR GXL polymerase and 9.4μl of distilled sterile water (TAKARA R050A). The following pairs of primers were used for genotyping Nodal^-/-^ line: (fwd TGAGGGTGAGAGGTGGGTG rev CTGCTGGATCGGAACTCAGG) and Nodal^ΔASE/ΔASE^ line (fwd AATTGTTTCTCCGTGGGCAG rev: AGCATCCCACTGATTTCCCA) Genotyping confirmed homozygous deletions.

### Production of Micropatterned adhesive substrates

The micro-patterned substrates used in the present study were produced using standard microcontact printing approach, presented in details in (Vedula et al., 2014). First, a mold with the desired pattern was produced by photo-lithography of SU8 resin on a silicon wafer. Uncured PDMS was then poured on these molds to obtain a negative replica of the SU8 master, after curing at 65C for at least 1h. Cured PDMS was then peeled of the SU8 master and inked with a solution of fibronectin (20ug/ml in PBS, incubation 30 minutes). The inked stamps were then washed twice with PBS and once with water and allowed to dry for 5 minutes. During that time, PDMS-coated glass cover slips (obtained by spin coating PDMS on glass and curing) were activated in a UV-Ozone cleaner. The Fibronectin coated stamp and the activated coverslip were then put into contact, for a few seconds and separated. To prevent cell adhesion outside of the desired areas, stamped coverslips were then incubated in PBS + 1% Pluronics F127 for at least 30 minutes. After 3 washes with PBS, the micropatterned substrates were ready for cell seeding or could be stored at 4C for a future experiment.

### EpiLCs differentiation on micropatterns

ESC cultures in N2B27+2i+LIF were harvested by trypsinization and seeded cells were first cultured for 24h on plates coated with fibronectin (15ug/ml for 30 min) in N2B27 supplemented with 1% KSR [life technologies, cat#10828010] and optionally with 20ng/ml ACTIVIN [Cell Guidance Systems] and 12ng/ml FGF2 [Cell Guidance Systems]. After 25h, cells were then dissociated using trypsin and seeded on micropatterned substrates at a density of 8000 cells per mm^2^, this high seeding density ensured to achieve a dense and uniform surface coverage. After 1h, cells that did not attach were removed by gentle flushing and changing the culture medium and remaining cells were allowed to form colonies for an additional 24h in 250μl of the same base medium. The medium was renewed every 24h during the course of the experiment. Colonies were stimulated by adding different combinations of factors to the medium (50ng/ml BMP4 [R&D System], 200ng/ml WNT3A [R&D System], 20ng/ml ACTIVIN and 12ng/ml FGF2, [cell guidance systems]). The stimulation was renewed after 24h, and colonies were cultured again in base medium.

### EpiSC culture and differentiation on micropatterns

EpiSC are derived from an E5.5 embryo and can be maintained in the primed state of pluripotency when cultivated in a medium containing Activin and FGF (Tesar et al., 2007). As previously reported (Sugimoto et al., 2015), we observed that it was necessary to add a Wnt secretion inhibitor (IWP2) to the EpiSC medium to prevent spontaneous appearance of small clusters of cells with anterior primitive streak identity, which prevented reproducible spatial organization of micropatterned colonies, see Fig. S1.

We thus maintained EpiSCs in Activin+FGF+IWP2 for at least 3 passages before seeding them on micro-patterned substrates. 24h hours after seeding on micropatterned substrates, IWP2 was replaced by BMP4 and spatial organization of the colonies was assayed 48h and 72h later by immuno-fluorescence.

### Immunostaining and microscopy

Cells were fixed with PBS + 4% paraformaldehyde [comp] at room temperature (RT) for 30 min, then washed twice with PBS for 10 min at RT, and blocked in PBS containing 0.1% Triton-X100 (PBST) and 3% bovine serum albumin for 1h at RT. They were then incubated overnight with the primary antibodies at 1:250 in the blocking solution (PBST+3% BSA). The cells were then washed 2 times 10 min in PBST and once for 40 min with the blocking solution, incubated for 2 hours with a cocktail of alexa fluorophore-coupled donkey anti-rabbit, goat, and mouse secondary antibodies at 1:500 [comp] and DAPI [conc, comp], wash three times in PBS (quick wash followed by 10 min and 50 min washes), and mounted in Prolong Anti-fade mounting medium [comp]. Fluorescent images of the colonies were acquired with a olympusIX81 inverted microscope equiped with a Yokogawa CSU-X1 spinning disk head. Radial profiles of fluorescence were extracted from maximum intensity projection using a custom matlab script.

### Quantitative analysis of gene expression

Cells from one well, containing between 7 and 10 colonies, were first dissociated in 200ul of 1M guanidinium by pipeting multiple times until the dissociation of all aggregates and the resulting solution was frozen at -20°C until all samples were collected. Cell total RNA was then extracted using the RNANucleoSpin RNA extraction kit (Macherey-Nagel), and quantified using a Nanodrop. Between 25 and 250ng of total RNA were then reverse transcribed (RT) using the Superscript IV VILO reverse-transcriptase (Thermo Fisher Scientific) and random hexamers according to manufacturer’s instructions. Gene expression was quantified by quantitative PCR on a LightCycler 480 Instrument II (Roche) using 1:250 of the reverse-transcribed RNA sample mixed with a specific primer pair (1uM final each, sequence in supplemental table) and 2X KAPA SYBR FAST qPCR Master Mix (Roche) according to manufacturer’s conditions. For each experiment, duplicates samples were analyzed. For each gene, the quality of the amplification was tested by quantifying its expression in a serially-diluted pool of all RT (1:10 to 1:320) to quantify amplification efficiency and target mRNA concentration relative to the RT pool in each sample.

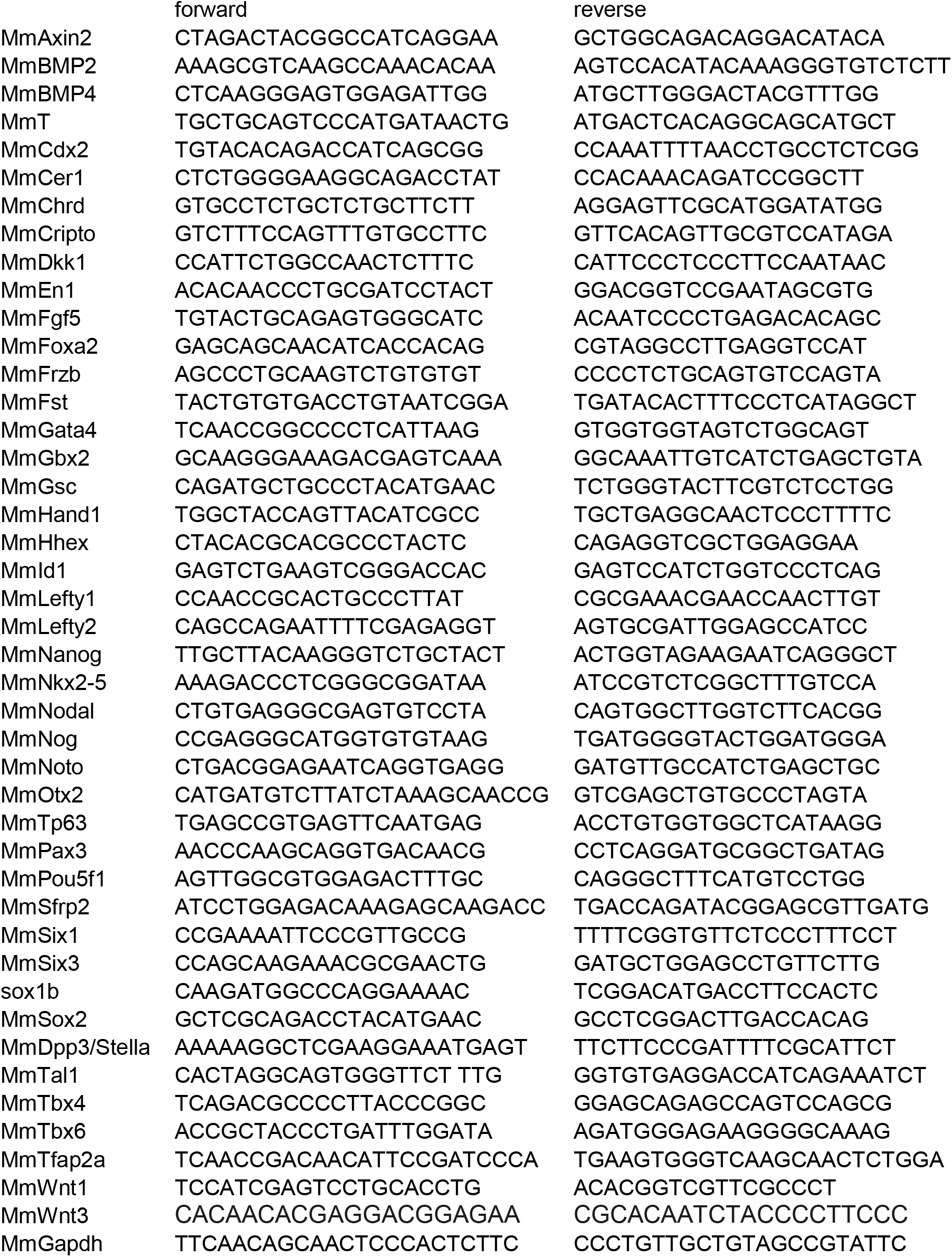
Table of PCR primer pairs used in this study

Table of antibodies used in this study

**Table of PCR primer pairs used in this study**
| **antigen** | **species** | **supplier** | **cat#** | **dilution** |
| --- | --- | --- | --- | --- |
| OCT3/4 | Mouse | BD Biosciences | 611203 | 1:250 |
| SOX2 | Rabbit | Abcam | ab97959 | 1:250 |
| NANOG | goat | r&D systems | AF2729 | 1:250 |
| BRACHYURY/TBX2 | Goat | R&D Systems | AF2085 | 1:250 |
| FOXA2 | Rabbit | Cell Signaling | 8186S | 1:250 |
| SOX17 | Goat | R&D Systems | AF1924 | 1:250 |
| P63 | Rabbit | Abcam | ab124762 | 1:250 |
| SOX1 | Goat | R&D Systems | AF3369 | 1:250 |
| CDX2 | Mouse | Abcam | ab157524 | 1:250 |
| E-CAD | Rabbit | Cell Signaling | 3195S | 1:250 |
| **secondary: donkey anti:** |  |  |  |  |
| **target species** | **fluorophore** | **supplier** | **ref** | **dilution** |
| Mouse | Alexa488 | Abcam | ab150109 | 1:500 |
| Rabbit | Alexa555 | Abcam | ab150062 | 1:500 |
| Goat | Alexa555 | Abcam | ab150134 | 1:500 |
| Goat | Alexa555 | Abcam | ab150134 | 1:500 |
| Mouse | Alexa647 | Abcam | ab150111 | 1:500 |
| Goat | Alexa647 | Abcam | ab150135 | 1:500 |
| Rabbit | Alexa647 | Abcam | ab150063 | 1:500 |

